# ELYS coordinates NF-κB pathway dynamics during development in *Drosophila*

**DOI:** 10.1101/386607

**Authors:** Saurabh Jayesh Kumar Mehta, Vimlesh Kumar, Ram Kumar Mishra

## Abstract

ELYS, a nucleoporin spatiotemporally regulates NF-κB pathway dynamics during development in *Drosophila* and its misregulation in post-embryonic stages leads to apoptosis mediated abnormalities.

**Abstract:** Nuclear pores are the exclusive conduit to facilitate the nucleocytoplasmic transport in a precisely regulated manner. ELYS, a constituent protein of nuclear pores, initiates assembly of nuclear pore complexes (NPCs) into functional nuclear pores towards the end of mitosis. Using cellular, molecular and genetic tools, here, we report that ELYS orthologue (*d*Elys) plays critical roles during *Drosophila* development. Through *in silico* analyses, we find all conserved structural features in *d*Elys except for the presence of non-canonical AT-hook motif strongly binding with DNA. *d*Elys localized to nuclear rim in interphase cells, but during mitosis, it was present on chromatin. RNAi mediated depletion of *d*Elys leads to aberrant development and defects in the nuclear lamina and NPCs assembly at the cellular level. Furthermore, we demonstrate that in *d*Elys depletion NF-κB is activated and accumulates inside the nucleus which results in illimed expression of critical molecules. *d*Elys depletion sustains NF-κB into the nucleus in post-embryonic stages. Prolonged NF-κB inside nucleus induces apoptosis in response to hitherto unknown quality check mechanism and highlights on the under-appreciated apoptotic paradigm of NF-κB pathway.

## Introduction

Nuclear pore complexes (NPCs) are a multi-protein assembly of nucleoporins (Nups), and their size varies from ~60 MDa in yeast to ~125 MDa in vertebrates. Nups are distributed into sub-complexes located on cytoplasmic, nuclear membrane and nucleoplasmic faces of nuclear pores (Hetzer, 2010; Hoelz et al., 2011). In metazoans, however, at the onset of mitosis, sub-complexes of NPCs dissociate from each other and gets redistributed inside the cell. The largest sub-complex of NPC called Y-shaped complex (hereafter Nup107 complex) has nine stoichiometric members (Ori et al.,2013). Nup107 complex plays important roles in NPC assembly, nucleocytoplasmic transport, and gets distributed to the kinetochores during mitosis (Orjalo et al., 2006;Zuccolo et al., 2007). ELYS (<u>E</u>mbryonic <u>L</u>arge molecule derived from <u>Y</u>olk <u>S</u>ac) was characterized in the mouse as a putative transcription factor important for hematopoiesis (Kimura et al., 2002). ELYS, also known as Mel-28, is required for the maintenance of the nuclear morphology and embryonic development in *C. elegans* (Fernandez and Piano, 2006; Galy et al., 2006). Although in yeast, ELYS is a small protein and carries the minimal ELYS domain, in the higher eukaryotes, its molecular mass ranges from ~190–250 kDa and is also known as AT-hook containing transcription factor 1 (ATCHF1). Mouse ELYS was characterized to contain nuclear localization signal (NLS), nuclear export signal (NES), N-terminal β-propeller region, central helical and most important, C-terminally located AT-hook motifs (Kimura et al., 2002; Okita et al., 2003). The β-propeller like domain present in many nucleoporins mediates interaction with other nucleoporins and facilitates NPC assembly (Bilokapic and Schwartz, 2013). In a breakthrough observation, ELYS was reported to be an integral member of the Nup107 complex (Franz et al., 2007; Rasala et al., 2006; Rasala et al., 2008). In interphase, ELYS is present at nuclear envelope and nucleoplasm, but in mitosis, it associates with chromatin, kinetochores, and spindles (Rasala et al., 2006). Importantly the conserved ELYS domain is required for its NPC and kinetochore localization (Gomez-Saldivar et al., 2016) where ELYS helps in microtubule polymerization (Yokoyama et al., 2014). ELYS tethers Nup107 complex to kinetochores and initiates their assembly into post-mitotic nuclear pores (Rasala et al., 2008). Loss of ELYS in *C. elegans* causes abnormal nuclear morphology and aberrant chromosome segregation that confers lethality (Fernandez and Piano, 2006; Galy et al., 2006).

ELYS is critically required for embryonic development as ELYS null mice die during embryonic stage E3.5 to E5.5, well before the onset of embryonic hematopoiesis (Kimura et al., 2002; Okita et al., 2004). However, conditional inactivation of ELYS locus in adult mice showed reduced effects and mice behaved normally (Gao et al., 2011). ELYS orthologue in Zebrafish, *flotte lotte (flo)*, is critically required for early embryonic and pharyngeal skeleton development (Davuluri et al., 2008). Additionally, *flo* deletion disrupted NPC formation and nuclear import induced replication stress in intestinal progenitors cells (de Jong-Curtain et al., 2009).

In addition to, playing an important role in NPC assembly, reduction in ELYS activity in HeLa cells leads to phosphorylation-dependent mislocalization of Lamin B Receptor (LBR) from the nuclear periphery (Mimura et al., 2016). ELYS recruits Protein phosphatase-1 (PP1) to the kinetochores and reduction in LBR phosphorylation levels counters the action of mitotic kinases (Hattersley et al., 2016). AT-hook motifs present in ELYS allow DNA binding and the trans-activation domains (acidic region) help in transcription (Kimura et al., 2002). AT-hook motif is rich in positively charged amino acids and harbors-RGRP-residues at its core, is sufficient for binding to DNA (Rasala et al., 2008). ELYS is also known to associate with the promoter region of actively transcribing genes (Pascual-Garcia et al., 2017). Thus, being a DNA binding protein, an effector of proper cell division and a regulator of nucleocytoplasmic trafficking, ELYS may have a significant role to play in early developmental events of an organism. However, no directed study has been performed on ELYS to obtain the mechanistic knowhow of important roles played by ELYS in cellular homeostasis and early developmental events.

We chose *Drosophila melanogaster* (fruit flies) to address the importance of ELYS in early developmental processes. Although CG14215 has been reported as ELYS in *Drosophila* (hereafter as *d*Elys) and shown to be present at the nuclear periphery and in association with promoter region on polytene chromosomes, none of these studies characterized *d*Elys for its role in NPC assembly or other cellular processes *in vivo* (Ilyin et al., 2017; Pascual-Garcia et al., 2017). Here, through genetic, molecular and cellular characterization we report that the *d*Elys plays important roles in the assembly and dynamics of nuclear pore and nuclear lamina functions. *d*Elys can bind with DNA through its non-canonical AT-hook motif, and RNAi mediated *d*Elys depletion induces severe lack of nucleoporins like Nup98, Nup43 and mAb414 in nuclear pore and perturbs lamin B receptor and lamin B and C incorporation into the nuclear lamina. Importantly, we observe that in *d*Elys depletion, developmentally important molecule Dorsal is activated and accumulated inside the nucleus and subsequently induces caspases (Dcp-1) to follow the apoptotic pathway. Our observations strongly imply that *d*Elys is a key developmental molecule required in cellular processes integral to normal cellular homeostasis. In addition to maintaining the nuclear architecture, *d*Elys contributes to the development by regulation of the apoptotic arm of dorsal pathway activation.

## Results

### ELYS is highly conserved in *Drosophila*

ELYS is a DNA binding nucleoporin which coordinates NPC assembly, and NPC mediated cellular processes like nucleo-cytoplasmic transport. Loss of ELYS induces developmental defects but how ELYS regulated events affect the complex process of development remains elusive. We have used *Drosophila* to address these questions. Although the CG14215 was reported to be *Drosophila* ELYS orthologue (*d*Elys), however, any analysis to attribute structural elements in the molecule and mechanistic details of ELYS function was absent. Primary sequence alignment of CG14215 exhibits overall ~20-21% identity with mouse and human ELYS (Fig. S1A). ELYS, being an AT-hook containing transcription factor, we first searched for the presence of AT-hook motifs in CG14215. By utilizing mouse ELYS AT-hook motif (Mm_Elys_AT-hook), we have predicted three AT-hook motifs, but importantly we noticed an evident lack of glycine and proline amino acids in the conserved-RGRP-core (overscored) present in several of AT-hook proteins (Fig.1A) (Aravind and Landsman, 1998). We asked if such a non-canonical AT-hook can bind chromatin. A carboxy-terminal fragment (aa 1858-1962) of *d*Elys bearing all three predicted AT-hook motifs, *d*Elys_AT-1, 2, and 3 helped determine the DNA binding abilities in Electrophoretic gel mobility shift assay (EMSA) (Fig. S1C). *d*Elys AT-hook bound with AT-repeat rich sperm dynein intermediate chain *(sdic)* promoter oligo in a dose-dependent manner and induced a clear shift in mobility (Fig.S1B). Our data further established that the presence of glycine and proline is not as important in the-RGRP-core, which is further supported by (Metcalf and Wassarman, 2006). Accordingly, we mutated the arginines in each predicted AT-hook motifs individually (mAT-1, mAT-2, and mAT-3) or combined mutations of arginines to alanines for all three AT-hooks (mAT-1+2+3) (Fig.S1C) and tested the ability of purified proteins (Fig.S1D) to bind with DNA. In these experiments, instead of oligo, we have used larger DNA fragment (~900bp) having an abundance of AT sequence (AT-rich) or a DNA fragment lacking them (Non-AT rich DNA). It is evident from the observations that both mAT-1and mAT-3 show a significant reduction in their ability to bind with AT-rich DNA (Fig.1B). However, the mAT-3 displayed pronounced perturbation in DNA binding abilities. Our mutational analysis of *d*Elys AT-hook motifs indicated that the AT-3 motif is the prominent DNA binding motif present at the C-terminus of the *d*Elys. Like other two predicted AT-hook motifs, AT-3 lacked glycine and proline in the core, but importantly it had alternate arginine residues as seen in canonical AT-hook of mouse ELYS. This particular feature made AT-3 the most probable AT-hook motif of *d*Elys. Moreover, our DNA binding observation also suggests that the AT-hook motif can serve as a general DNA binding protein also (Fig.S1E). Quantification of the DNA binding data showed that AT-hook motif-1 and 3 are the DNA binding motifs with AT-3 having a stronger affinity for AT-rich DNA sequence (Fig.1C and Fig. S1F).

**Figure 1:**
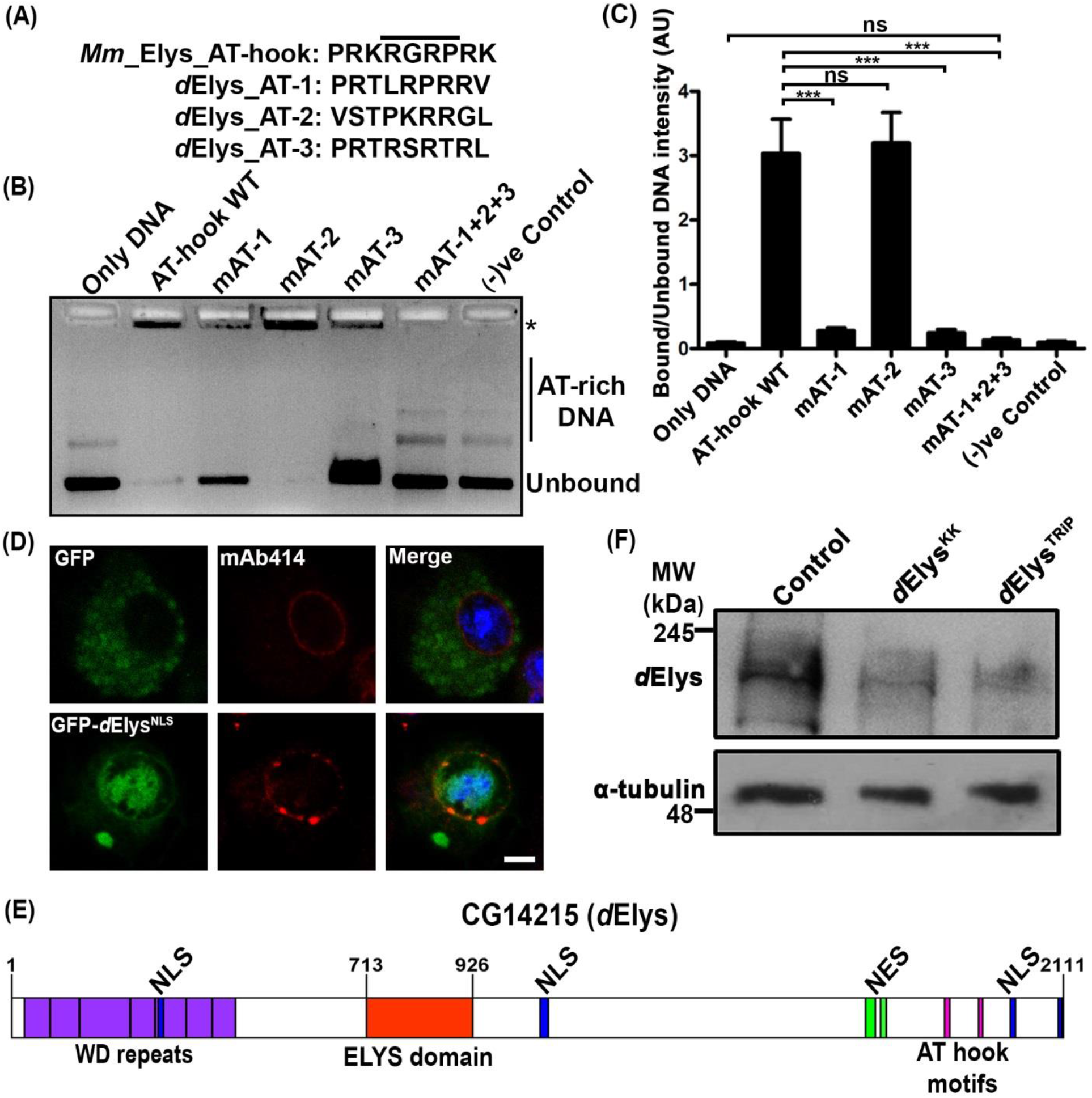
ELYS is structurally and functionally conserved in *Drosophila:* (A) Three predicted AT-hook DNA binding motifs of CG14215 are compared with canonical mouse AT-hook motif. Core RGRP signature residues of AT-hook are marked. (B) DNA binding ability of predicted AT-hooks is tested in EMSA experiment. AT-rich DNA binding by purified AT-hook fragment (AA. 1858-1962) and mutated AT-hook proteins. Type of protein used is indicated above each lane. Asterisk indicates bound DNA with decreased mobility. (C) Quantification of DNA binding ability of each AT-hook fragment used. Quantification is derived from at least three independent experiments. *** represents p<0.0001, ns is non-significant. Error bar represents a Standard error of the mean (SEM). Statistical analysis is derived from one-way ANOVA followed by post-hoc Tukey’s multiple comparison tests. (D) Nuclear localization of GFP (in green) fused with N-terminal NLS of *d*Elys. Only GFP remains in the cytoplasm. Nuclear periphery is marked with mAb414 (in red). DNA is stained with DAPI. (Scale bar: 1 μm) (E) *d*Elys shows conserved protein motifs similar to mammalian ELYS. It consists of a WD repeat at the N-terminus (purple), an ELYS domain in the middle (orange) and an AT-hook motif at the C-terminus (pink). It also contains nuclear import and export signal motifs (blue and green respectively). The N-terminal NLS has been used for functional analysis. (F) An antibody generated against *d*Elys identifies a band of ~235 kDa in *Drosophila* third instar larva head lysate in control. RNAi mediated depletion of *d*Elys is reflected as a decrease in band intensity. α-tubulin was used as loading control.

Mammalian ELYS was shown to have a strong nuclear localization signal (NLS) and nuclear export signal (NES) sequences. Our *in silico* predictions indicate for the presence of multiple NLS spread throughout the molecule and two C-terminal NES in **d*Elys*. When the N-terminal NLS of *d*Elys (strongest among the predicted NLS) was fused with GFP and expressed in *Drosophila* S2 cells, it was effectively transported inside the nucleus indicating its functionality (Fig.1D).

Further predictions of structural features present in *d*Elys find a conserved arrangement of N-terminal β-propeller domain (aa 1-488), central helical domain (aa 489-977) and a C-terminal unstructured region (Fig. S1A). Homology-based 3D structure predictions on the N-terminal β-propeller domain of *d*Elys, identified a highly conserved and significantly overlapping seven-bladed beta-propeller structure as seen in the N-terminal domain of mouse ELYS (Fig. S1G) (Bilokapic and Schwartz, 2013; Kelley et al., 2015). Moreover, we could also predict for the presence of WD repeats at the N-terminal region and an ELYS domain in the central helical region of the *d*Elys (Fig. 1E). Evolutionary divergence analysis indicates that *d*Elys is present in a clade quite distinct and distant from vertebrate ELYS (Fig. S1H). Our analysis strongly showed that *d*Elys carries all the conserved structural motifs present in canonical ELYS despite low primary sequence homology.

To address the *d*Elys functions in cellular and developmental processes in greater detail, we have raised polyclonal antibodies against the C-terminal fragment (aa 1796–2111). Affinity purified antibodies recognized ~230 kDa band in *Drosophila* salivary gland lysate (Fig.1E) as well as in lysates obtained from *Drosophila* S2 cells (Fig.S2A). We also observed that this antibody could detect a concurrent decrease in *d*Elys protein level when *d*Elys is silenced using RNAi, indicating strongly towards specificity of *d*Elys detection by the antibody. In tissues, *d*Elys antibodies detected specific and strong nuclear rim staining along with feeble cytoplasmic annulate lamellae staining in early *Drosophila* syncytial embryos (Fig. 2A and Fig.S2C). This particular observation highlighted about the relatively low abundance of *d*Elys in annulate lamellae as compared to other nucleoporins detected by mAb414 (Fig. S2E). *d*Elys antibodies observed a pattern at nuclear rim overlapping with GFP-Nup107 which is a known interacting partner of ELYS across eukaryotes and also with the nuclear lamina-associated Lamin-B receptor in salivary gland nuclei (Fig. S2B, D).

**Figure 2:**
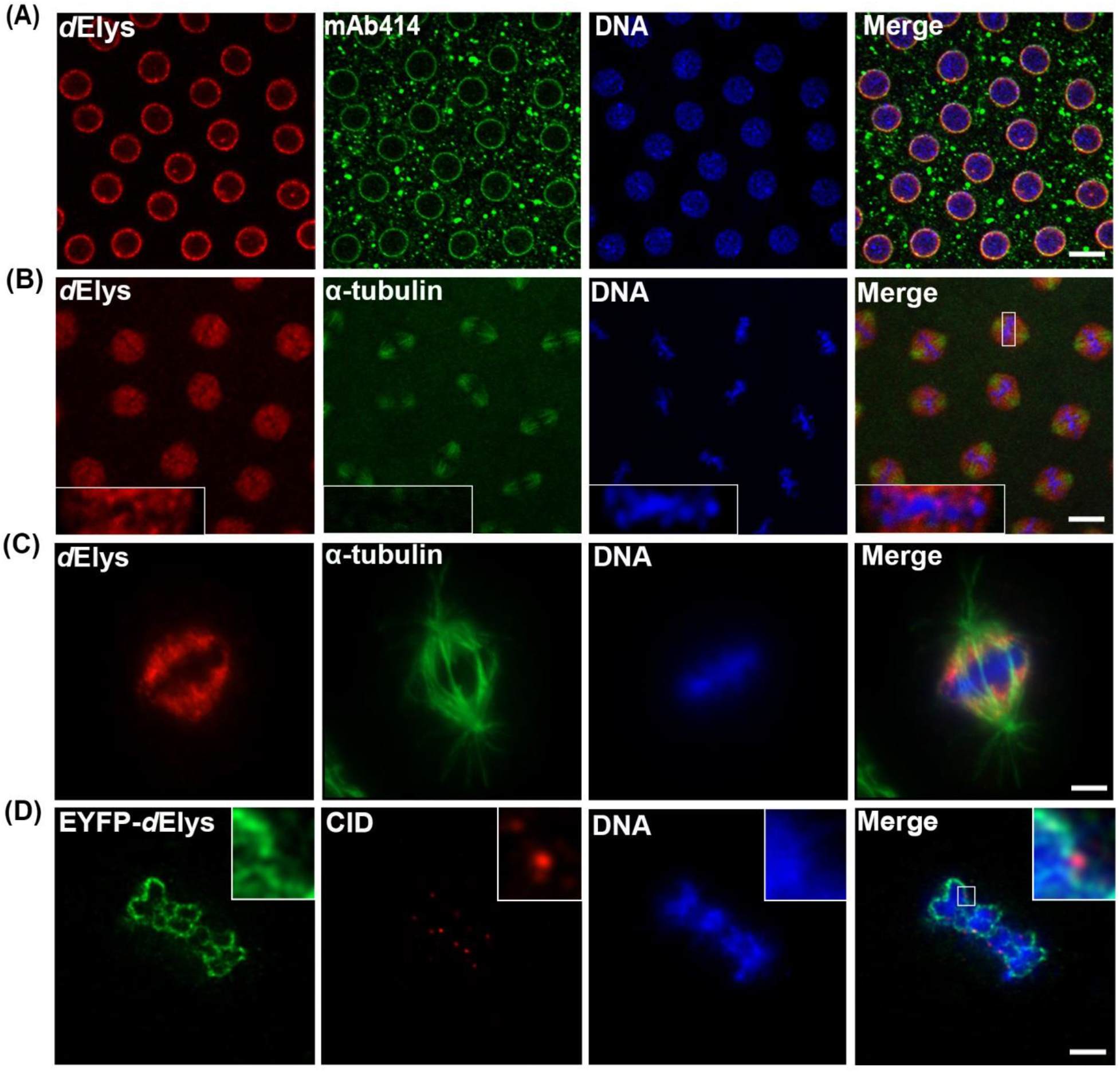
*d*Elys is present at nuclear envelope during interphase but translocate in the vicinity of kinetochore during mitosis. (A) Conserved localization of *d*Elys on a nuclear envelope during interphase identified by the *d*Elys antibody (red) in syncytial *Drosophila* embryos. mAb414 (green) stained nuclear pore complexes and DAPI stained DNA. (Scale bar: 5 μm) (B) Mitotic syncytial embryo showing localization of *d*Elys (red) in spindle envelope area, partial co-localization with α-tubulin (green) is evident. Inset showing *d*Elys localization around mitotic chromosomes. (Scale bar: 5 μm, Inset: 0.5 μm) (C) Metaphase arrested *Drosophila* S2 cells highlighting localization of *d*Elys (red) around mitotic chromosome in a cup-shaped manner. (Scale bar: 2 μm) (D) A high-resolution image of metaphase-arrested *Drosophila* S2 cells showing localization of EYFP tagged *d*Elys (green) around the mitotic chromosome. Kinetochores are stained with anti-centromere identifier (CID) antibody (red). (Scale bar: 2 μm)

### *d*Elys is absent from kinetochores and localizes around chromosome in mitosis

ELYS exhibits cell-cycle dependent variations in its localization and is present at the nuclear rim during interphase and to the kinetochores in mitosis (Franz et al., 2007;Rasala et al., 2006; Yokoyama et al., 2014). We probed if *d*Elys also exhibits a cell-cycle dependent sub-cellular localization change. The nuclei in *Drosophila* early embryos divide synchronously and grow in syncytium which provides the desired window to observe cell-cycle dependent localization of proteins. As observed earlier, *d*Elys was present at the nuclear rim during interphase syncytial embryo (Fig. 2A), but in mitosis, a diffused staining is obvious in the entire presumptive nucleoplasmic region of syncytial embryos (Fig. 2B, Fig. S2E). However, a careful analysis of higher magnification images, indicate that *d*Elys is present around the mitotic chromosomes (Fig. 2B inset). Interestingly, *d*Elys staining resembled a bracket to chromosomes in *Drosophila* S2 cells arrested in mitosis, where is showed partial co-localization with a-tubulin spindle fibres (Fig. 2C). We further asked if similar to other reported ELYS like molecules, *d*Elys is present at kinetochores in addition to a chromatin bracket in mitosis. Co-staining of kinetochores with *Drosophila* CENP-A homolog (CID) (Vermaak et al., 2002) in EYFP-*d*Elys expressing S2 cells exhibited a distinct absence of *d*Elys staining from kinetochores. *d*Elys did not co-localize with CID but were in close vicinity of it (Fig. 2D). The EYFP-*d*Elys, when expressed in salivary glands, produced a localization pattern similar to endogenous *d*Elys (Fig. S2F). It is important to note that ELYS recruits the Nup107 complex to the kinetochores in cell culture, *Xenopus* egg extracts, and *C. elegans* embryos. However, the Nup107 complex was reported to be absent from kinetochores in *Drosophila* syncytial mitotic embryo (Katsani et al., 2008). Our observation that *d*Elys is absent from kinetochores provides a suitable explanation for lack of Nup107 complex staining from *Drosophila* kinetochores.

### *d*Elys is essential for normal development in *Drosophila*

After characterizing as a nucleoporin, we asked whether *d*Elys is essential for development in *Drosophila*. Over-expression of *d*Elys using UAS-*d*Elys transgenic line did not yield any observable phenotypic differences when compared with wild-type (data not shown). *d*Elys was depleted *in vivo* using RNAi line (v103547) from Vienna *Drosophila* Resource Centre (VDRC)(*d*Elys^KK^) to understand its role in cellular functioning. We also generated TRiP based RNAi line (*d*Elys^TRiP^) to validate phenotypes observed by *d*Elys^KK^ RNAi line, which do not have any predicted off-targets. Ubiquitous knockdown of *d*Elys in *Drosophila* using *Act*5C-GAL4 had severe consequences on viability with variable lethality at each developmental stages, no adults emerged from one of the RNAi line *d*Elys^KK^, and only a few viable flies (~4%) emerged out in *d*Elys^TRiP^ (Fig. 3A). We observed that *d*Elys depleted embryos are showing ~4% lethality while ~46% lethality occurring at the larval stage, ~50% of pupae died in the late pupal stage of development (Fig. 3A). The extent of *d*Elys depletion in RNAi lines when assessed by quantitative PCR showed that *d*Elys^KK^ RNAi line caused ~70% knockdown, while *d*Elys^TRiP^ leads to ~55% knockdown in *d*Elys transcript levels (Fig. 3B). Although *d*Elys^KK^ has two predicted off-targets in addition to *d*Elys, we have not observed a significant change in the level of off-targets (Fig.S3). As the *d*Elys^KK^ line is showing maximum knockdown of *d*Elys, we have used *d*Elys^KK^ RNAi line for all subsequent studies. *d*Elys has been contributed in high amount maternally, which makes it difficult to analyze its role in early stages of development. To overcome this problem, we used *mat-a-tub-* GAL4 which depletes maternally contributed *d*Elys before induction of zygotic transcription. Maternally depleted *d*Elys (*d*Elys^KK^ RNAi line) embryos have an abnormal shape as well as they do not show pole cells development (Fig. 3C, arrowhead). Hatching rate analysis showed that maternal *d*Elys depleted embryos showed ~62% lethality compared to control which showed ~12% lethality (Fig.3D).

**Figure 3:**
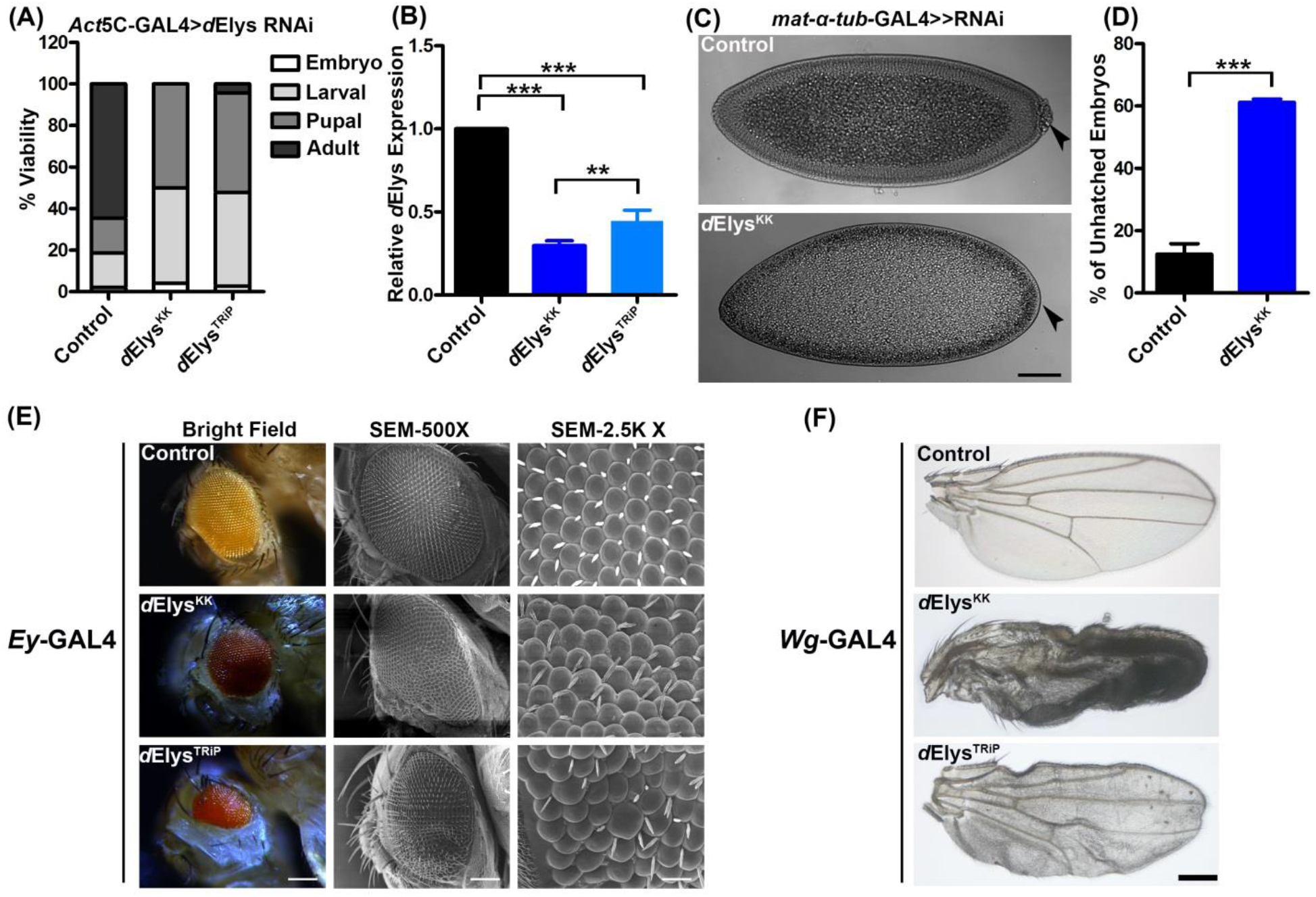
*d*Elys is essential for normal development of *Drosophila*. (A) Quantification representing lethality stages of the *d*Elys knockdown organism. *d*Elys was knockdown using two independent RNAi lines driven with *Act*5C-GAL4. (B) Quantitative PCR for *d*Elys knockdown using two different RNAi lines. Data represented from at least three independent experiments. Statistical significance derived from one way ANOVA followed by Tukey’s post-hoc test. Error bar represents SEM. *** represents p<0.0001 and ** represents p<0.001. (C) Early embryonic knockdown of maternal *d*Elys using *mat-a-tub-GAL4* showed defects in embryo development. Arrow-head indicates pole cells. (D) Hatching rate analysis of maternal *d*Elys depleted embryos compared to control. (E) Eye-specific knockdown of *d*Elys using the Ey-GAL4 drivers. Column 1 shows the stereomicroscopic image, column 2 represents the SEM image (500 X) and column 3 shows an SEM image (2.5K X). Observations were made from at least three independent experiments. (Scale bar: 200 μm in the stereomicroscopic image, 20 μm in SEM column 1 and 2 μm in SEM column 3) (F) *d*Elys depletion in wing using the Wg-GAL4 driver. Representative images are shown for wing phenotypes. Observations were made from at least three independent experiments. (Scale bar: 200 μm)

To avail more insights on the importance of *d*Elys in development, we silenced *d*Elys in a tissue-specific manner with *d*Elys depletion in eyes and wings using eyeless (Ey) and wingless (*Wg*) drivers respectively. Eyes in *d*Elys depleted flies appeared shrunk with a significant reduction in the area occupied by ommatidia in addition to irregular size and disarrangement of ommatidia leaving behind eye cavity. We also observed that when *d*Elys was depleted significant number of ommatidia show loss of eye bristles, duplication of bristles and their orientation was irregular (Fig. 3E). Similarly, *d*Elys silenced wings look crumpled with signs of vein atrophy and significant cell death in wing blade tissues (Fig. 3F). The severity of these phenotypes varied between the two RNAi lines targeting *d*Elys yet the observations made regarding eye and wing phenotypes were consistent. Our data revealed that the presence of *d*Elys is important for the normal development of *Drosophila*.

### Loss of *d*Elys results in the nuclear pore and nuclear lamina assembly defects

We proposed that the developmental defect observed in *d*Elys knockdown organisms would be the most probable outcome of compromised NPC assembly. To analyze if *d*Elys is important in NPC assembly, we assessed the localization of various nucleoporins representing various sub-complexes of NPC in the *Drosophila* salivary gland nuclei. Nup43, a stable member of the Nup107 complex, Nup98, a mobile nucleoporin and mAb414 reacting to nucleoporins with FG repeats and regulate cargo movement through NPCs were absent from the nuclear membrane and presented a diffused cytoplasmic staining in *d*Elys depleted salivary gland tissue (Fig.4A-D). Lack of nucleoporin sub-complexes and lack of functional nuclear pores in *d*Elys silenced nuclei reinforces the idea that *d*Elys is a critical player in functional NPC assembly. Concomitantly, we observed a marked increase in cytoplasmic staining for these nucleoporins in *d*Elys depleted salivary gland tissues highlighting possible increase in annulate lamellae (AL) structures in the cytoplasm. Quantitation of these intensities also confirms our observation of loss of these nucleoporins from the nuclear periphery (Fig.4E-H). We have checked the protein levels of these nucleoporins in larval head-complex lysate by western blotting, and the observations suggest no significant decrease in the protein levels of tested nucleoporins except for Nup98 (Fig.4Q). Since DAPI intensities didn’t change between control and *d*Elys silenced tissues; we have used to normalize the variability observed with the intensity of each molecule tested. We have further confirmed with Histone H3 staining that DNA intensities remain unaltered upon *d*Elys depletion (Fig. S4). Our data with *d*Elys further strengthens the conserved role for ELYS in NPC assembly throughout metazoans. We argued that if *d*Elys depletion perturbs NPC assembly, then it must compromise the nucleo-cytoplasmic transport as well. *In situ* detection of mRNA showed that *d*Elys depleted salivary gland nuclei had more mRNA compared to control nuclei (Fig.S6 A and B), highlighting defect in nuclear export. When we expressed GFP-tagged SV-40 NLS under ubiquitin promoter and the nuclear import of GFP-NLS was severely compromised upon *d*Elys depletion while it normally localized to the nucleus under control depletion conditions (Fig. S6C and D). We could detect a faint GFP-NLS signal in the nucleus under *d*Elys silenced condition which is possible due to residual NPCs present in the nuclear membrane or maybe because of interphase NPC assembly that occurs independently of *d*Elys presence. These observations emphasized important roles played by *d*Elys in NPC mediated nucleo-cytoplasmic transport of vital molecules.

**Figure 4:**
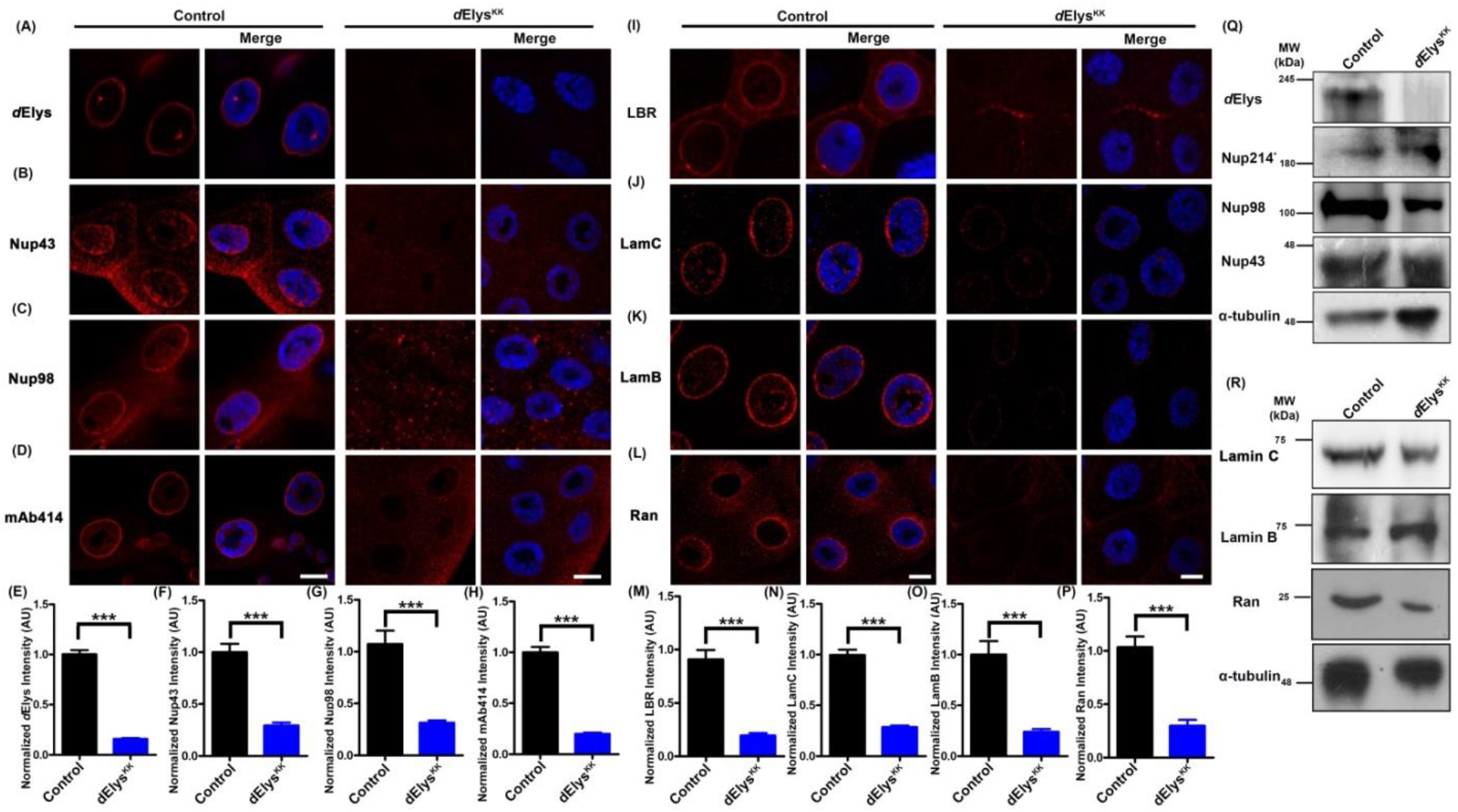
*d*Elys is essential for nuclear pore complex and nuclear lamina assembly in *Drosophila*. (A-D) Localization of *d*Elys, Nup43, Nup98 and FG-repeat nucleoporins (detected by mAb414) on the nuclear rim assessed in control and *d*Elys depleted (*Act*5C-GAL4 driven) salivary gland nuclei. Indicated molecules are represented in red. DNA is stained with DAPI. (Scale bar: 5 μm) (E-H) Quantitation of signal intensities of *d*Elys, Nup43, Nup98 and mAb414 on nuclear rim compared to control. Intensities of each molecule were normalized against the intensity of DAPI. Data is represented from at least three independent experiments. Error bar represents SEM. Statistical significance derived from student’s t-test. *** represents p<0.0001. (I-K) Localization of LBR, Lamin C, Lamin B which constitute nuclear lamina was assessed on the nuclear rim in control and *d*Elys depleted (*Act*5C-GAL4 driven) salivary gland nuclei. Each indicated molecules is represented in red. DNA is stained with DAPI. (Scale bar: 5 μm) (L) Nuclear transport factor Ran is analyzed in control and *d*Elys down-regulated (*Act*5C-GAL4 driven) nuclei. (Scale bar: 5 μm) (M-P) Quantitation of nuclear rim localization of each indicated molecules in control and *d*Elys RNAi. Intensities of each molecule are normalized against the intensity of DAPI. Data is represented from at least three independent experiments. Statistical significance derived from student’s t-test. Error bar represents SEM. *** represents p<0.0001, ns is non-significant. (Q, R) Western blot analysis of each molecule tested in control and *d*Elys knockdown (*Act*5C-GAL4 driven) third instar larva head lysate. * indicates detected with mAb414. Molecular markers are indicated next to the blot. α-tubulin was used as a loading control.

ELYS plays critical roles in NPC assembly and also interacts with LBR. Consequently, ELYS regulates nuclear lamina organization and incorporation of LBR in nuclear envelope in a phosphorylation-dependent manner (Clever et al., 2012; Mimura et al., 2016). We promptly investigated the changes occurring in nuclear lamina architecture upon *d*Elys depletion. Salivary gland nuclei from *d*Elys silenced organism showed reduced localization of Lamin B-receptor (LBR), Lamin B and Lamin C in the nuclear lamina (Fig.4I-K). Quantitation of nuclear rim intensities for these molecules further highlights the significant decrease in recruitment of these molecules upon *d*Elys depletion (Fig.4M-O). Our observation is the first ever report highlighting the importance of *d*Elys in the localization of Lamin B and Lamin C to nuclear lamina. We also observe that recruitment of *d*Elys to the nuclear envelope is independent of LBR as the *d*Elys staining pattern remained unaffected in RNAi mediated depletion of LBR (Fig. S5C-E). These observations suggested that *d*Elys works independent and upstream of LBR. *d*Elys helps in the incorporation of LBR, Lamin B, and C into the nuclear lamina which is indispensible for the maintenance of the nuclear lamina and ultimately the nuclear architecture.

Nucleo-cytoplasmic transport of protein cargo through NPC is coordinated by Ran-GTPase which can be found on the nuclear rim as well. Importantly we have noticed that *d*Elys depleted salivary gland tissues show striking lack of Ran reactivity from nuclear periphery and its redistribution was seen throughout the cell (Fig. 4L, P). Lack of Ran-GTPase from nuclear periphery may also be one of the reasons for defective nucleo-cytoplasmic transport in *d*Elys silenced condition. The absence of Ran-GTPase in addition to lack of Nups and lamins prompted us to ask if everything associated with nuclear periphery observes re-distribution upon *d*Elys knockdown. However, when we probed for nuclear periphery associated TATA-box binding protein-1 (TBP1, dTBP in *Drosophila*)(Choudhury et al., 2017) localization in *d*Elys depleted salivary gland tissues, it was unperturbed and marked characteristic nuclear periphery staining pattern (Fig. S5A, B). To probe changes in the protein level of nuclear lamina molecules upon *d*Elys knockdown, we performed western blotting on larval head complex lysate and with antibodies corresponding to these molecules. Lamin B and Lamin C protein level were unaffected in *d*Elys knockdown while Ran showed little decrease in *d*Elys knockdown (Fig. 4R). These observations approve the important role for *d*Elys in the nuclear pore and nuclear lamina assembly. Consequently, loss of *d*Elys could contribute to a severe reduction in the nuclear transport of various developmentally important molecules leading to abnormal development.

### *d*Elys knockdown shows activated NF-κB pathway and induced apoptosis

To deduce the underlying molecular mechanism that drive *d*Elys depletion induced defects, we sought to check for perturbed nucleo-cytoplasmic traffic of various signaling molecules. We observed significant nuclear accumulation of key developmental transcription factor, Dorsal (NF-κB) under *d*Elys depleted conditions in salivary gland tissue. Under normal condition, Dorsal stably associates with cactus and is rendered inactive in the cytoplasm but rapidly shuttles inside the nucleus when activated in response to certain stimuli (Rushlow et al., 1989). Shuttling of dorsal inside nucleus is accompanied by ubiquitination-dependent degradation of the inhibitor of NF-κB (IκB) in the cytoplasm. (Belvin et al., 1995; Whalen and Steward, 1993). We depleted *d*Elys using RNAi, and the assessed Dorsal signal in salivary gland tissues. In control tissues Dorsal signal was diffused and was present prominently in the cytoplasm. However, the *d*Elys depleted cells show intense dorsal signal in close association with chromatin inside the nucleus (Fig. 5A). Quantification of the dorsal signal from *d*Elys RNAi tissues shows a marked increase in nuclear dorsal signal (Fig. 5D). Nuclear localization of Dorsal indicates loss of its cytoplasmic binding partner Cactus, which freed Dorsal. To ensure this, we examined the cytoplasmic level of cactus and observed that cytoplasmic cactus signal decreased significantly in *d*Elys depletion when compared to control tissues (Fig. 5B). This loss of cactus staining was quantified, and the data corroborated the cactus degradation and dorsal activation (Fig. 5E). Our data strongly suggest that *d*Elys silencing trigger the activation of NF-κB pathway. Dorsal activation and movement to the nucleus is a proliferative signal and developmentally important event. However, we observe cell death which is a probable consequence of the unfavorable accumulation of dorsal inside the nucleus. Different studies suggest that activated dorsal can also induce apoptosis in cells (Kaltschmidt et al., 2000; Radhakrishnan and Kamalakaran, 2006; Ryan et al., 2000).

**Figure 5:**
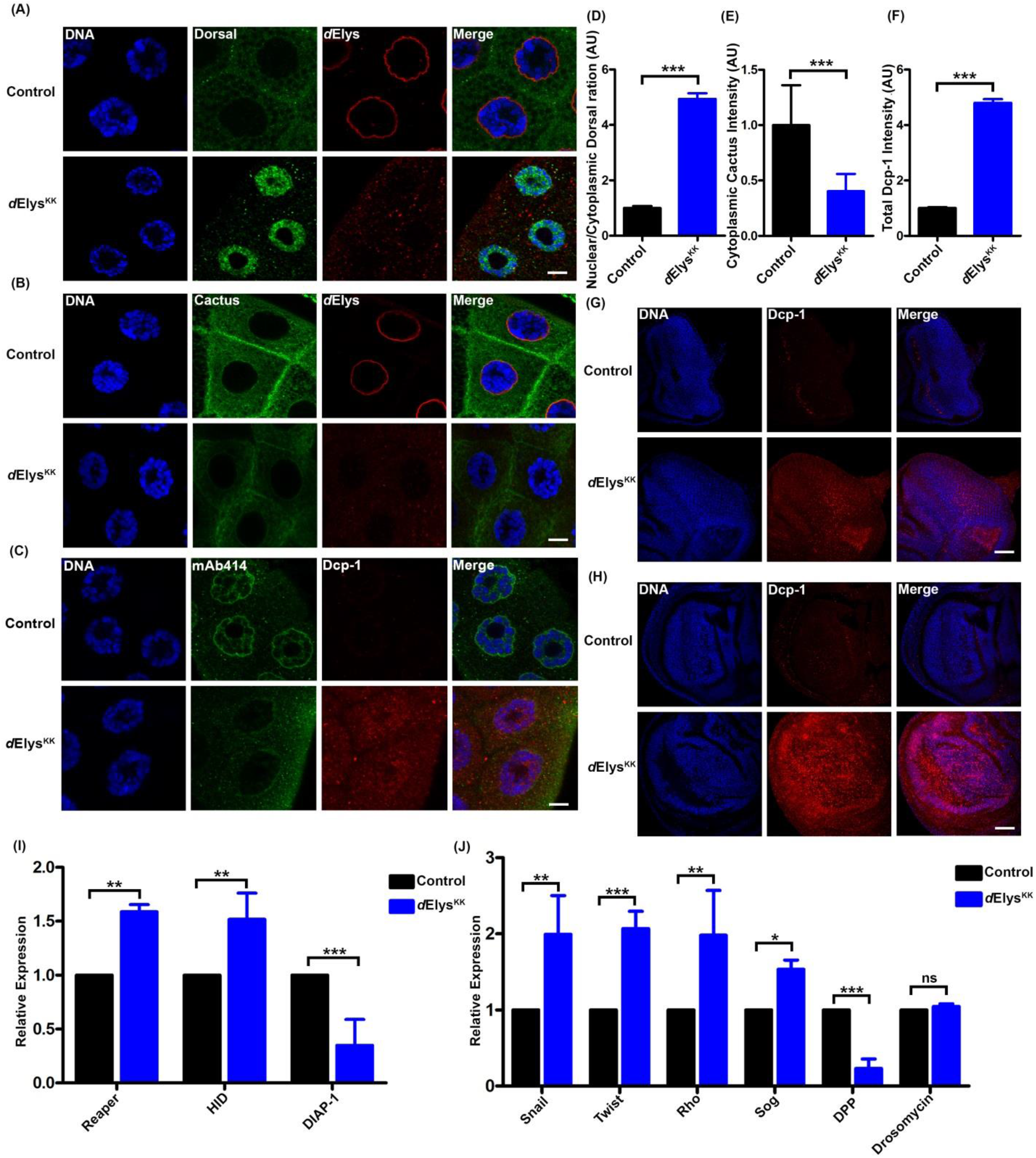
NF*-k*B is activated in *d*Elys depletion and show induced apoptotic response. (A) Nuclear localization of Dorsal was assessed with anti-Dorsal (green) in control and *d*Elys RNAi (*Act*5C-GAL4 driven) salivary gland nuclei. DNA is stained with DAPI. (Scale bar: 5 μm) (B) Cactus was detected in the cytoplasm with anti-cactus (green) in control and *d*Elys depleted (*Act*5C-GAL4 driven) salivary gland cells. DNA is stained with DAPI. (Scale bar: 5 μm) (C) Apoptotic induction in salivary gland cells was analyzed using the anti-Death caspase-1 antibody (red) in control and *d*Elys depletion (*Act*5C-GAL4 driven). NPCs are marked with mAb414 (green). DNA is stained with DAPI. (Scale bar: 5 μm) (D-F) Quantification of intensities of indicated molecules in control and *d*Elys RNAi condition. The intensity of each molecule was normalized to the intensity of DAPI. Data is represented from at least three independent experiments. Statistical significance derived from student’s t-test. Error bar represents SEM. *** represents p<0.0001. (G) *Drosophila* third instar eye imaginal discs from control and *d*Elys knockdown (*Act*5C-GAL4 driven) larva were assessed for apoptotic induction using Dcp-1 (red). DNA is stained with DAPI. (Scale bar: 20 μm) (H) Wing imaginal discs from control and *d*Elys depleted (*Act*5C-GAL4 driven) larva were assessed for apoptosis using Dcp-1 (red). DNA is stained with DAPI. (Scale bar: 20 μm) (I) Apoptosis regulator genes Reaper, *Drosophila* inhibitor of Apoptosis-1 (DIAP-1), and Hid were analyzed by quantitative PCR in control and *d*Elys RNAi (*Act*5C-GAL4 driven). Data is represented from at least three independent experiments. Statistical significance derived from students test. Error bar represents standard deviation. *** represents p<0.001 and ** represents p<0.01. (J) NF-κB target genes Snail, Twist, Rho, decapentaplegic (DPP), Short gastrulation (Sog) were analyzed by quantitative PCR in control and *d*Elys RNAi (*Act*5C-GAL4 driven). An immune response target of NF-κB, Drosomycin was tested for its expression. Data is represented from at least three independent experiments. Statistical significance derived from student’s t-test. Error bar represents standard deviation. *** represents p<0.001 and ** represents p<0.01 * represents p<0.05, ns is non-significant.

We next probed if the abnormal development of eye and wing tissues as well as lethality observed upon *d*Elys depletion could be a result of increased cell death via apoptosis. We used acridine orange staining in salivary glands as a primary assay to infer the induction of apoptosis. *d*Elys depleted tissues accumulated significantly more acridine orange than normal tissues which suggested activated apoptosis (Fig. S6 E, F). To further assert that *d*Elys depletion indces apoptotic response, we probed for *Drosophila* caspase-1 (Dcp-1) which is critical for normal embryonic development and elevation in Dcp-1 levels is a hallmark for apoptosis (DeVorkin et al., 2014; McCall and Steller, 1998; Song et al., 1997). Dcp-1 antibodies staining showed significantly enhanced reactivity for Dcp-1 and significantly more number of Dcp-1 positive puncta were obvious in *d*Elys silenced salivary gland tissues (Fig. 5C, F). Since the *d*Elys depletion induced significant defects in eye and wing development, we asked if the apoptosis sets in early during the development of these organs in the absence of *d*Elys. We probed for the Dcp-1 levels in Wing and eye imaginal discs obtained from wild-type and *d*Elys depleted third instar larva. Both the eye and wing imaginal discs showed increased punctuate reactivity with Dcp-1 upon *d*Elys depletion. (Fig. 5G, H). We next asked if NF-κB causes any alterations in the expression of apoptotic genes. We probed this by quantitative real-time PCR for pro-apoptotic genes Reaper and Hid as well as for anti-apoptotic DIAP-1(Drosophila Inhibitor of Apoptosis-1) transcripts in cDNA prepared from the third instar larval head complex of control and *d*Elys depleted organism. We observed a significant increase in the level of pro-apoptotic Reaper and Hid expression while there was a significant decrease in the level of anti-apoptotic DIAP-1 upon *d*Elys depletion (Fig. 6I). The relative levels of DIAP-1, Reaper, and Hid are important determinants of cell survival or cell death. Our qPCR observations and Dcp-1 immunostaining data exhibit undisputable correlation and indicate towards apoptotic induction upon *d*Elysdepletion. Dorsal, being a transcription factor and activated in the nucleus in *d*Elys knockdown should bring a change in the levels of its target genes. To investigate this, we checked the levels of primary target genes of Dorsal in cDNA prepared from the third instar larval head complex by quantitative PCR. While relative levels of Snail, Twist, Rho and Sog observed an increase, the levels of Dpp (Decapentaplegic), a gene suppressed by dorsal, decreased in *d*Elys depleted samples as compared to control (Fig.5J). To rule out the possibility of activated NF-κB pathway under immune response, we checked for the expression of the anti-microbial peptide, Drosomycin in *d*Elys depleted cDNA. We found that the expression of Drosomycin is unaltered, suggesting that activation of NF-κB pathway does not have a microbial infection and immunity component to it (Fig.5J).

**Figure 6:**
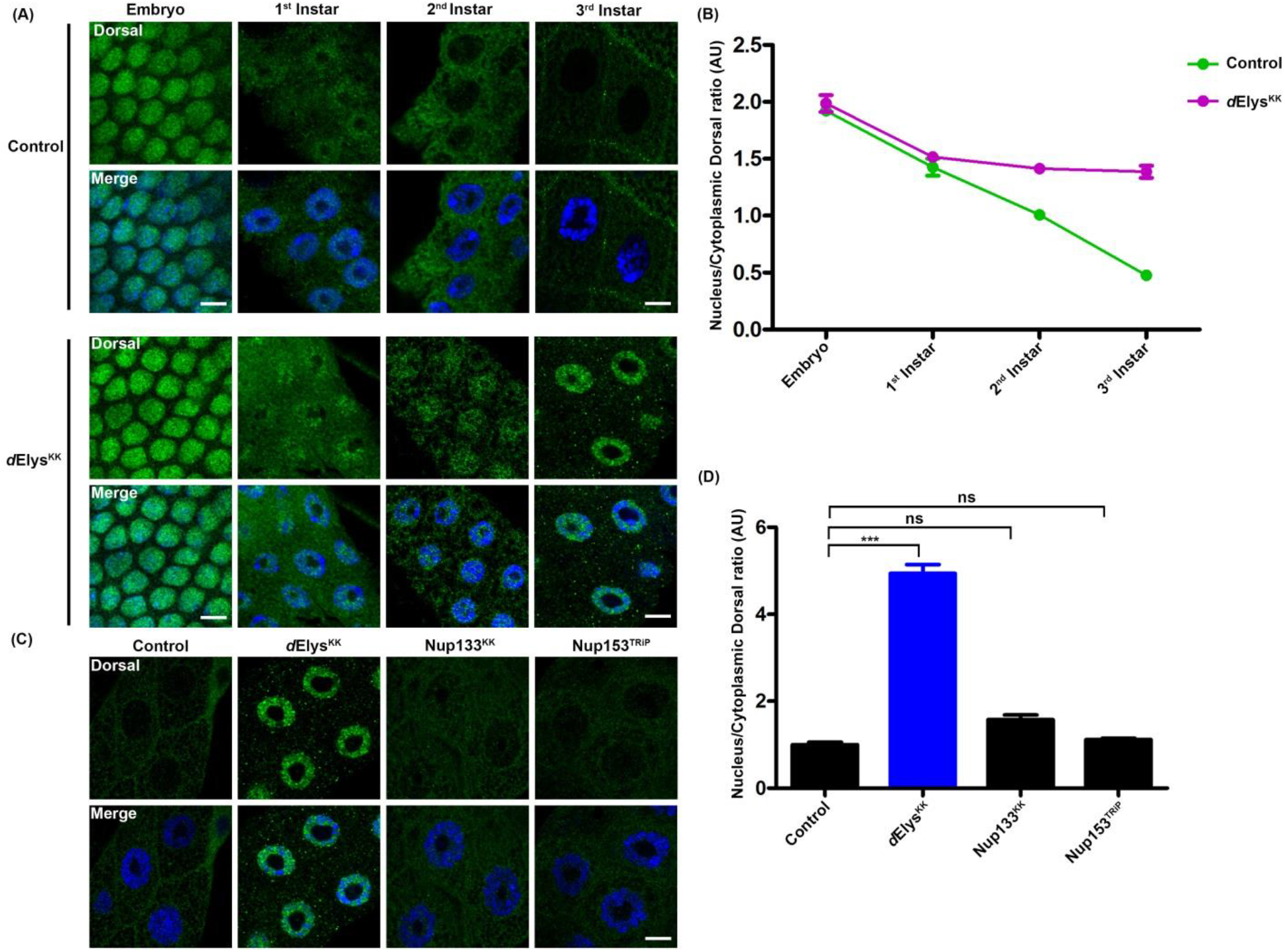
*d*Elys depletion stabilizes Dorsal in the nucleus during development and specific to *d*Elys. (A) Localization of Dorsal (green) assessed in control and *d*Elys depletion in early stages of development. *d*Elys depleted (*Act*5C-GAL4 driven) nuclei showed increase localization of Dorsal in the nucleus compared to control. DNA is stained with DAPI. (Scale bar: 5 μm) (B) Quantitation of nuclear/cytoplasmic intensity ratio of Dorsal measured through developmental stages of *Drosophila* showing sustained localization of Dorsal in the nucleus in *d*Elys depletion. Data is represented from at least three independent experiments. Error bar represents SEM. (C) Dorsal nuclear localization assessed in control and nucleoporin knockdown salivary glands (*Act*5C-GAL4 driven). DNA is stained with DAPI. (Scale bar: 5 μm) (D) Quantitation of nuclear/cytoplasmic intensity ratio of dorsal in control and nucleoporins knockdown salivary gland nuclei. (*Act*5C-GAL4 driven). Data is represented from at least three independent experiments. Statistical significance derived from one way ANOVA followed by Tukey’s post-hoc test. Error bar is SEM. *** represents p<0.0001 and * represents p<0.05, ns is non-significant.

Dorsal targets Snail, Twist, Rho, Dpp, and Sog expresses early in development. We used third instar larval head complex for cDNA preparation, which is quite late in the development and still detected the expression of these molecules. Sustained expression of these molecules till late developmental stages might be the cause of apoptosis in *d*Elys depletion which leads to developmental defects. Snail, Twist, Dpp, and Rho are already shown to be activators of the apoptotic response in *Drosophila* (Campbell et al., 2018; Gullaud et al., 2003; Lim and Tomlinson, 2006; Neisch et al., 2010). It is thus not surprising to find sustained expression of these molecules till late developmental stages resulting into induction of apoptosis asserting developmental defects in *d*Elys depletion conditions.

### Dorsal distribution to the nucleus is sustained in post-embryonic tissue and specific to *d*Elys nucleoporin depletion

Dorsal is one of the maternally contributed molecules which is required for early developmental progress (Belvin et al., 1995; Govind, 1999). Dorsal nuclear localization is affected in all ventral nuclei of developing embryos specifying dorsoventral polarity. During early development, Dorsal after getting phosphorylated keeps shuttling between cytoplasm and nucleus under the influence of upstream stimuli (DeLotto et al., 2007; Perkins, 2006). The nuclear presence of Dorsal gradually decreases with progress in development beyond the embryonic stage particularly after zygotic induction and Dorsal remains in the cytoplasm through stable association with Cactus. We set out to investigate if nuclear localization of Dorsal a temporal event of later developmental stages or Dorsal entrapped in the nucleus even after the embryonic stage in the *d*Elys knockdown organism. Firstly, we analyzed the dorsal levels in embryonic stages and noticed that there is no observable difference in the nuclear signal of Dorsal in *d*Elys depleted and control embryos (Fig. 6A, First vertical panels). We then subsequently looked for dorsal signals in salivary gland nuclei isolated from each successive larval stages. Similarly, no apparent difference was seen in the dorsal signals inside the nuclei isolated from first instar control and *d*Elys depleted larval salivary glands (Fig.6A, second vertical panels). However, we noticed a gradual but steady decrease in nuclear Dorsal signal in embryonic and 1^st^ instar stage nuclei of control and *d*Elys depletion conditions. While, *d*Elys depleted salivary gland nuclei continued to have more Dorsal in 2^nd^ and 3^rd^ larval stages, control nuclei has only residual or no Dorsal (Fig.6A, third-fourth vertical panels). The sustained nuclear localization of Dorsal becomes apparent in the 3^rd^ instar larval salivary stage. (Fig.6A, third vertical panel). This observation finds a complementary decrease in cytoplasmic signal of Dorsal in the *d*Elys depleted salivary glands. Quantification of nuclear Dorsal signal intensities from control and *d*Elys depleted salivary gland nuclei further establish that Dorsal is retained in the nucleus during post-embryonic developmental stages upon *d*Elys depletion (Fig.6B). Our results strongly suggest that *d*Elys plays important role in shutdown of activated NF-κB signaling during post-embryonic stages and its sequestration in the cytoplasm.

We next asked if this sustained nuclear localization of Dorsal is *d*Elys specific or a consequence of lack of nuclear pore complexes upon *d*Elys depletion. To check this, we investigated dorsal localization in salivary gland nuclei of critical nucleoporins Nup160, Nup133, Nup107, and Nup153. Members of the Nup107 complex play a key regulatory role in post-mitotic and interphase NPC assembly while Nup153 is a critical molecule for interphase NPC assembly only (Orjalo et al., 2006; Vollmer et al., 2015; Zuccolo et al.,2007). When compared with control depletion Nup153 and Nup133 depletion show no significant difference in Dorsal signals inside nuclei as in control (Fig.6C). Although, Nup160 and Nup107 depletion cause lethality at the 1^st^ instar larval stage, the Dorsal intensities inside the nucleus were indistinguishable from those observed in the nuclei of 1^st^ instar larvae of wildtype organism (Fig. S7A). Quantitation of nuclear intensities of the Dorsal signal demonstrate that Dorsal signal inside nucleus increase by ~5 fold when *d*Elys is depleted but the same remains largely unchanged upon depletion of other nucleoporins involved in NPC assembly (Fig.6D and Fig.S7B). This highlights that sustained residence of Dorsal inside is *d*Elys specific and not the outcome of the absence of NPCs in nuclear envelop.

### *d*Elys depletion-induced apoptosis is Dorsal mediated

We next examined if apoptosis and nuclear localization of dorsal is an independent consequence of *d*Elys depletion. We have created a genetic combination of *d*Elys depletion in the Dorsal null background. However, this imparted lethality at 1^st^ instar larval stage hence was not useful. We decided to combine the knockdown of *d*Elys and Dorsal and analyzed the apoptotic response from co-depleted salivary glands. *d*Elys depletion induced accumulation of dorsal and enhanced punctate nuclear staining of Dcp-1, but the co-depletion of *d*Elys and Dorsal showed a reduction in Dcp-1 and dorsal staining in salivary gland nuclei (Fig.7A). Quantitation of signal intensities further indicated that Dcp-1 signals return to normal when *d*Elys and Dorsal are co-depleted (Fig. 7B, C). This also highlighted the fact that induction of apoptosis is dorsal dependent. Moreover, RNAi line used for Dorsal knockdown does not deplete maternally contributed Dorsal and thus do not affect embryonic development.

**Figure 7:**
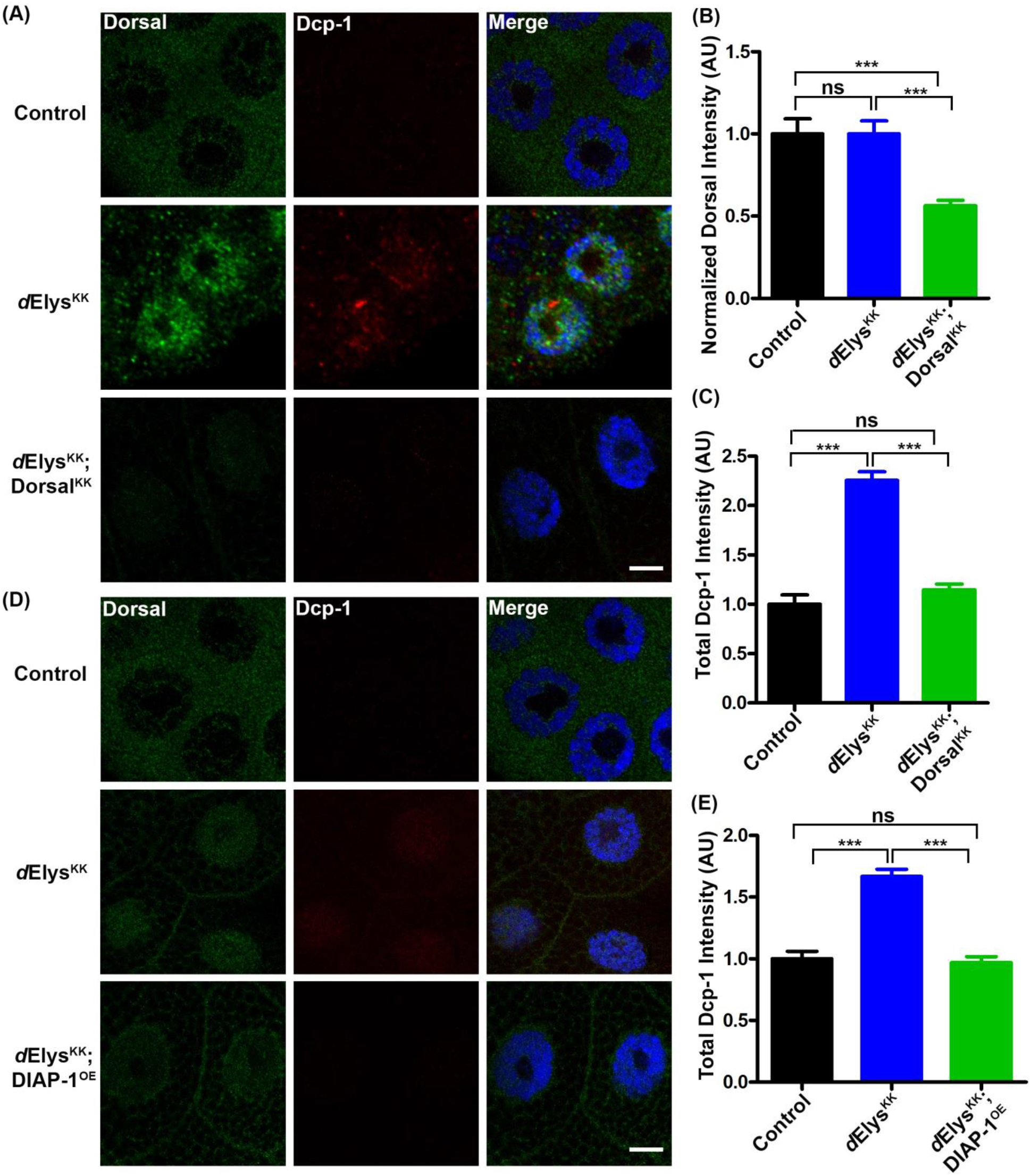
Dorsal depletion and DIAP-1 overexpression can rescue *d*Elys depletion effects. (A) Apoptotic induction assessed in control, *d*Elys RNAi and *d*Elys, Dorsal double knockdown (*Act*5C-GAL4 driven). Salivary gland cells were stained with Dorsal (green) and Dcp-1 (red). DNA is stained with DAPI. (Scale bar: 5 μm) (B, C, and E) Quantitation of signal intensity of Dorsal and Dcp-1 in indicated samples. Data is derived from at least three independent experiments. Statistical significance derived from one way ANOVA followed by Tukey’s post-hoc test. Error bar represents SEM. *** represents p<0.0001 and ns is non-significant. (D) Apoptotic induction assessed in control, *d*Elys RNAi and *d*Elys RNAi; DIAP-1 over-expressed (Salivary gland-specific expression) salivary gland cells. Cells stained with Dorsal (green) and Dcp-1 (red). DNA is stained with DAPI. (Scale bar: 5 μm)

We next decide to examine if apoptosis can be rescued by overexpressing DIAP-1 in the *d*Elys knockdown. As ubiquitous overexpression of DIAP-1 is lethal, we used localized expression using tissue specific drivers. The salivary gland-specific *d*Elys depletion elicited the increased Dcp-1 signal inside the nucleus, but the DIAP-1 overexpression in *d*Elys depletion background brings the Dcp-1 levels back to normal and subsiding the apoptotic response (Fig.7B). Although the visualized intensity of Dcp-1 is different as compared to Fig.7A, Quantitation of Dcp-1 intensities verified that inhibition of apoptotic response was a consequence of DIAP-1 over-expression in *d*Elys depletion background (Fig.7E). Our data strongly suggest that the induction of apoptosis upon *d*Elys depletion is Dorsal mediated and the DIAP-1 over-expression counters the *d*Elys depletion-induced apoptosis.

**Figure 8:**
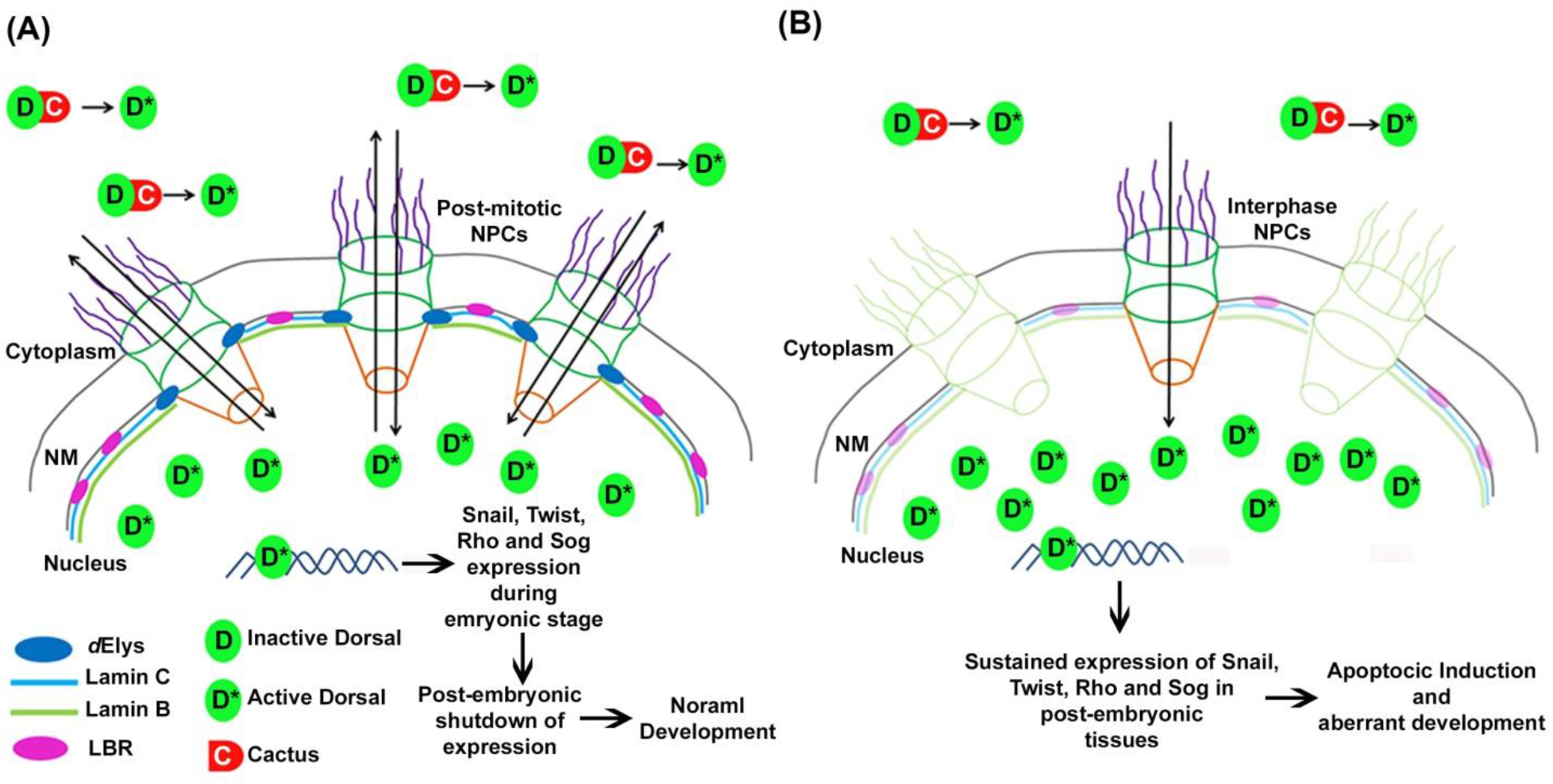
The Model highlighting role of *d*Elys in Dorsal mediated development. (A) *d*Elys initiates postmitotic NPC assembly leading to functional nuclear pore complexes and the nuclear lamina in the nuclear membrane. Developmental nuclear-cytoplasmic shuttling of Dorsal occurs normally leading to normal development. Dorsal transferred back to the cytoplasm after completing its transcription factor role in the nucleus by some post-translational modifications or may be degraded (DeLotto et al.,2007). (B) Depletion of *d*Elys causes mislocalization of other nucleoporins and NPC assembly is perturbed. The nuclear lamina is also defective in a *d*Elys knockdown. Dorsal entered in the nucleus during early developmental stages remains in the nucleus in *d*Elys depletion may be due to the defective post-translational processing of Dorsal in the nucleus. Sustained Dorsal presence in nucleus activates mistimed transcription of Dorsal target genes which needs to be shut post-embryonic stage. This mistimed transcription may lead to apoptosis and ultimately lethality and developmental defects. Thus *d*Elys presence is critical for the fate of Dorsal localization and ultimately to the development of an organism.

## Discussion

ELYS was identified as a putative transcription factor in mouse embryos and was also found in association with chromatin (Gillespie et al., 2007; Kimura et al., 2002). Subsequently, target genes for ELYS, and cellular function of ELYS remained elusive for quite some time. Studies on ELYS later gained attention when it was found in stable association with a Nup107 complex of nuclear pores (Franz et al., 2007; Galy et al., 2006; Rasala et al., 2006). Nuclear pores present a rim staining around chromatin in interphase cells, and Nup107 complex gets recruited to kinetochores in mitosis. Similarly, ELYS antibodies recognized specific nuclear rim reactivity in interphase cells and an overlapping kinetochore staining pattern was seen with Nup107 complex (Rasala et al., 2006). Later, it was established that ELYS recruits Nup107 complex to kinetochores and hence helps in regulation of Nup107 complex mediated microtubule formation at kinetochores (Gillespie et al., 2007; Mishra et al., 2010; Yokoyama et al.,2014). Importantly, ELYS binds with promoter regions in chromatin and interacts with Mcm-7 helicase to exert control during DNA replication as well as interacts with Piwi for transcriptional regulation (Gillespie et al., 2007; Ilyin et al., 2017). Loss of ELYS induces severe defects in organism development like skeleton loss in Zebrafish, embryonic lethality in mouse, tissue differentiation defects and aging phenotypes (Cerveny et al.,2010; Davuluri et al., 2008; de Jong-Curtain et al., 2009; Gao et al., 2011; Okita et al.,2004). Together, these observations suggested an important regulatory role for ELYS in organism homeostasis and development through a conserved underlying molecular mechanism.

Our study set out to address homeostatic roles for ELYS in *Drosophila* development. We report that the *Drosophila* ELYS orthologue (*d*Elys) encoded by CG14215 has conserved structural and functional elements of a canonical ELYS molecule. We demonstrate that *d*Elys can efficiently bind to AT-rich DNA despite possessing a non-canonical AT-hook motif (Fig. 1). Expectedly, *d*Elys showed nuclear rim staining in interphase which is characteristics of nucleoporins, however, in mitosis, it misses kinetochores and envelopes chromatin. High-resolution imaging for *d*Elys observes discrete chromatin periphery localization absent from the kinetochores in mitotic cells (Fig. 2). This result is in striking contrast with its higher orthologue which showed kinetochore localization during mitosis (Gillespie et al., 2007; Mishra et al., 2010;Yokoyama et al., 2014). This observation concurs with the lack of Nup107 complex from kinetochores reported in *Drosophila* and provides a suitable explanation for the same (Katsani et al., 2008).

RNAi mediated ubiquitous knockdown of *d*Elys, induced developmental defects and lethality in *Drosophila* (Fig. 3). The lethality, however became obvious during the later developmental stages (the 3^rd^ instar larval), and most probable explanation for this delay is the maternal contribution of *d*Elys in embryos. Therefore, *d*Elys depletion in eyes and wing tissues produced more pronounced morphological defects (Fig. 3).Further, the loss of nuclear pores severely affecting nuclear morphology, and nucleo-cytoplasmic transport probably is the reason for the observed lethality (Aitchison et al.,1995; Emtage et al., 1997; Fabre and Hurt, 1997; Fernandez and Piano, 2006; Galy et al., 2006). Moreover, we report a conserved interaction between *d*Elys and Lamin B receptor (LBR) which helps in the organization of the nuclear lamina (Clever et al.,2012; Mimura et al., 2016). Nuclear pore assembly after mitosis in *Drosophila* is a stage-wise process, and mature NPCs are seen largely in interphase. Importantly, different stages of NPC assembly become apparent during different stages of mitosis (Kiseleva et al., 2001). *d*Elys knockdown adversely affects recruitment of various Nups representing different sub-complexes of NPC inducing a gross loss in NPC morphology (Fig. 4). Importantly, we report a novel observation that in addition to LBR, both Lamin B and C are also missing from the nuclear lamina (Fig. 4). LBR interacts with lamin B, and thus the loss of LBR upon *d*Elys depletion may induce loss of lamins too. Moreover, the Ran-GTPase critically required for nucleo-cytoplasmic transport of protein cargoes shows redistribution from the nuclear periphery upon *d*Elys depletion (Fig.4). The Ran-GTPase mislocalization in association with lack of NPCs and compromised nuclear lamina cause the severe nuclear transport defects of important regulatory molecules. *d*Elys is thus critical for nuclear pores, nuclear lamina organization, and nuclear import/export and together these are integral to the maintenance of cellular equilibrium. Our search for developmentally important molecular players whose functionality is affected significantly and contributes to *d*Elys phenotypes narrowed down to Dorsal (NF-κB). Dorsal disengages from its inhibitor (IκB) and accumulates inside nucleus in its activated form (Whalen and Steward, 1993). Induction of Dorsal signaling plays an important role in the growth and development of the organism (Espin-Palazon and Traver, 2016; Govind, 1999). In *Drosophila,* Dorsal is present inside the nucleus during early developmental stages or when microbial infection elicits an immune response. We observe for the first time that Dorsal is present inside the nucleus even beyond the early development stages, and without any obvious infection which activates death caspase-1 (Dcp-1) (Fig.5). We thus demonstrate that NF-κB pathway activation upon *d*Elys depletion independent of microbial infection and even when any evidence of the antimicrobial peptide synthesis is lacking. We further showed that early development targets of Dorsal, namely Snail, Twist, Rho, Dpp, and Sog are expressed even during 3^rd^ instar larval stage (Fig.5). The persistent and undesired activation of developmental players until the later stages of development might activate the checkpoint alarming the system of a misprogramming upon *d*Elys depletion. The activated checkpoint can subsequently cease the development by inducing apoptosis. Our data corroborated this idea, and an increase in apoptosis was evident in *d*Elys depleted salivary gland tissues, eye and wing imaginal discs (Fig. 5). We attribute critical roles to *d*Elys in early development, and lack of *d*Elys leads to persistent accumulation of activated Dorsal inside nucleus even during post-embryonic developmental stages (Fig.6). ELYS is known to target protein phosphatases to the nucleus, which might dephosphorylate and exports molecules back to the cytoplasm (Hattersley et al., 2016). It is prudent to suggest that a similar phosphatase may be regulated by *d*Elys to control nucleo-cytoplasmic shuttling of Dorsal in *Drosophila*. It will be interesting to identify if such a phosphatase is involved in the dorsal shuttling process in *Drosophila*.

Our observation regarding Dorsal pathway activation is in direct contrast to what has was shown for another nucleoporin *mbo* (Nup88) in *Drosophila*. Depletion of *mbo* neither affected nuclear import-export nor nuclear lamina assembly. More importantly, nuclear levels of activated Dorsal were reduced in response to infection when Nup88 is deleted (Uv et al., 2000). Nup88 (Genenncher et al., 2016) along with other nucleoporins like Nup62 (Yang et al., 2015), Nup153 and Nup214 (Xu et al., 2002) are shown to be critical in developmental pathways, but detailed analyses emphasizing on their involvement in selective nuclear translocation of activated signals. In contrast, we report sustained activation of NF-κB signaling and its accumulation inside the nucleus only upon *d*Elys depletion but not with other nucleoporins involved nuclear pore complex assembly. Our results highlight the key difference between *d*Elys and other nucleoporins and affirm a role for *d*Elys in developmental signaling in addition to NPC and nuclear lamina organization.

Dorsal has lately been reported for inducing apoptotic responses, and emerging ideas suggest that Dorsal, depending on cues, helps in growth versus apoptotic fate choices (Barkett and Gilmore, 1999). Further the activation of the pro-apoptotic function of NF-κB is cell-type specific and signal-dependent (Silverman and Maniatis, 2001). The persistent activation and accumulation of NF-κB upon *d*Elys depletion could be a result of upstream signaling induced by yet an unidentified inducer. Our observations suggest a nuclear pore independent function of *d*Elys which impinges on pro-apoptotic functions of NF--κB. The apoptosis induction and lethality in imaginal discs and wing blade defects (cell death and atrophy) could be dorsal mediated apoptotic outcomes of *d*Elys loss. The prolonged nuclear accumulation of Dorsal shifts balance towards apoptosis causing tissue death and accompanying developmental defects (Fig. 5). Cells may be resorting to the NF-κB pathway dependent apoptosis as one of the ways to counter this defect and elimination of cells with abnormal nuclei produced upon *d*Elys depletion. Moreover, there may be additional signaling cascades that are perturbed by changes in *d*Elys levels, which requires further elucidation.

We suggest that *d*Elys levels help maintain a tight balance between pro-growth and pro-apoptotic responses to help achieve normal growth and development in *Drosophila*. *d*Elys does regulate development and could contribute to normal growth in post-mitotic cells by keeping the NF-κB trigger under control in a spatiotemporal manner (Fig. 6). We present *d*Elys levels as few examples which probably can activate NF-κB pathway by intracellular cues in contrast to much appreciated NF-κB pathway activation by extracellular stimuli for growth or infection (Gillespie and Wasserman, 1994; Silverman and Maniatis, 2001).

Our study is a novel report showing an intricate regulatory mechanism which cell relies on to ensure proper development in *Drosophila*. It also adds a new dimension in various roles played by the nucleoporin ELYS. *d*Elys depletion showed constant activation of NF-κB pathway in *Drosophila,* but how upstream signaling is continued its signals to this pathway, requires further detailed analysis. We speculate that various checkpoint like molecules present in the cell might be sensors for NPC and nuclear lamina assembly defects and activates regulatory pathways in accordance. We anticipate greater roles of nucleoporins in cellular homeostasis maintenance rather than being a mere structural component of NPC, which could be an intriguing question for further research.

## Materials and Methods

### *In silico* analysis

An orthologue of ELYS in *Drosophila* was identified by using mouse ELYS as the reference sequence. PSI-BLAST with three iterations using mouse ELYS and its AT-hook motif sequence was used against the non-redundant protein database at NCBI with default parameters. ELYS orthologues in other organisms were identified by similar search in the NCBI database. T-coffee, multiple sequence alignment server was used for alignment (Notredame et al., 2000). The aligned sequence was then used for visualization in Jalview and highlighted with clustal X color scheme. The phylogenetic tree was inferred from this analysis using Jalview (Waterhouse et al., 2009). Protein sequence motifs were identified using the SMART database for the representative sequence from each organism (Schultz et al., 1998). IBS was used to draw scaled protein illustration for each protein (Liu et al., 2015). Percentage identity was calculated against mouse ELYS as a reference for each ELYS like molecules. Secondary structure for CG14215 was predicted using PSIPRED v3.3, DisoPRED and DomPRED (Jones,1999). Predicted secondary structure for N-terminal and central helical domain were compared with the experimentally proven secondary structure of human and mouse ELYS(Bilokapic and Schwartz, 2013). 3D structure of NTD of CG1425 was modeled using a Phyre2 algorithm and mouse ELYS NTD structure as a template (Kelley et al.,2015).

### Fly strains and genetics

All flies were reared at 25°C on standard corn meal-yeast-agar medium. RNA interference crosses were grown at 28°C for better expression of GAL4. RNAi line (KK 103547) for CG14215 was obtained from the Vienna *Drosophila* Resource Centre (VDRC). Other fly lines used in this study were obtained from the Bloomington*Drosophila* Stock Centre (BDSC) at Indiana University. Controls used in this are F1 progeny from Driver line crossed with W^1118^ flies. Fly lines used in this study are mentioned in table S1. All the combination and recombination are made with standard fly genetics. Tissue-specific knockdown of *d*Elys was achieved by *Act*5C-GAL4, *Ey*-GAL4 and Wg-GAL4 drivers obtained from BDSC.

### Cloning and Transgenic fly generation

We obtained the clone LD14710 containing full-length CG14215 coding sequence in pBluescript SK (−) vector from *Drosophila* Genomics Resource Centre (DGRC). This clone was used as a template for cloning full-length CG14215 into pTVW (Vector. No. 1091), a fly transformation vector with gateway cassette, UASt promoter, and N-terminal EYFP tag as well as in pTW (Vector. No. 1129) vector containing gateway cassette and UASt promoter (obtained from DGRC gateway-1 collection). Cloning was done using Gateway cloning kit as per manufacturer’s instructions (Thermo Fisher Scientific). RNAi lines against CG14215 was generated by cloning 503 bps from exon 7 of CG14215 predicted using SnapDragon algorithm at *Drosophila* RNAi screening center (DRSC) website (FlyRNAi.org) which do not have any predicted off-targets (Flockhart et al.,2012) and cloned into gateway based pVALIUM10 RNAi vector obtained from DRSC (TRiP). Transgenic flies were generated at Fly facility of Centre for Cellular and Molecular Platforms at National Center for Biological Sciences (C-CAMP-NCBS), Bangalore, India. Primers used in this study are mentioned in table S2.

### *Drosophila* S2 cell culture and transfections

*Drosophila* S2 cells were grown in Schneider’s *Drosophila* medium (Gibco, Thermo Fisher Scientific) supplemented with 10% Foetal bovine serum (Gibco, Thermo Fisher Scientific) and 50 U/ml penicillin, 50 U/ml streptomycin and 25 μg/ml amphotericin B (Gibco, Thermo Fisher Scientific) at 25°C in CO_2_ free incubator. Full-length CG14215 was cloned in pAVW (Vector No. 1087) from DGRC. S2 cells were transfected at 50% confluency with Effectene transfection reagent (Quiagen). Cells were grown for 3 days post-transfection. S2 cells were arrested in metaphase of mitosis by addition of 20 μM MG-132 (RTU solution, Sigma) for 2 hours at 25°C.

### Antibody generation and western blotting

C-terminal 343 amino acid of CG14215 which is unique, most antigenic and hydrophilic part was amplified from LD14710 and cloned into pET28a (+). Protein was expressed in *E.coli* BL21 (DE3) cells, induced using 200 μM IPTG (Sigma) and incubated at 18°C overnight. Cells were pelleted down, lysed in 100 mM NaH_2_PO_4_, 10 mM Tris-HCl buffer, pH 8.0 containing 8 M urea and 1% Triton X-100, 50 μg/ml lysozyme and 1X protease inhibitor cocktail (Roche). Recombinant protein was purified over Ni-NTA beads and eluted with low pH at 4.5 in 100 mM NaH_2_PO_4_, 10 mM Tris-HCl buffer, pH 8.0 containing 8 M urea. Protein was dialyzed against 100 mM NaH_2_PO_4_, 10 mM Tris-HCl buffer, pH 8.0 containing decreasing amount of urea up to 2 M. Protein was concentrated using centrifugal concentrator of 30 kDa (Amersham, GE healthcare Life sciences). Protein was flash froze in liquid nitrogen and stored at −80°C. Polyclonal antibody against CG14215 was generated in the rabbit at Abgenex, Bhubaneswar, Odisha. Antibodies were affinity purified over purified antigen chemically cross-linked to N-hydroxysuccinimidyl-sepharose (NHS) beads (Sigma). Eluted with low pH, neutralized and dialyzed against PBS overnight at 4°C. Antibody against full-length ďNup43 was also generated using identical protocol.

Larval head complexes were dissected in cold PBS from wandering third Instar larva and lysed in lamellae buffer. Total 2 head complexes equivalent were loaded in each well of the gel. *Drosophila* S2 cells were pelleted down and lysed in 50 mM Tris-HCl pH 7.6, 150 mM NaCl, 1 mM MgCl_2_,1 mM EDTA, 10% glycerol,0.4% sodium deoxycholate, 1 % Triton X-100, 0.5% SDS, 1 % NP-40, 2X PIC (modified from (Andlauer et al.,2014). 5 and 10 μg of total protein was resolved on to 8% SDS-PAGE and transferred to methanol activated PVDF membrane (Merck-Millipore). Polyclonal, anti-*d*Elys antibody was used at 1:1000, anti-Nup43 antibody at 1:500, anti-Nup98 at 1:2500, anti-Lamin B at 1:2500, anti-Lamin C at 1:2500, anti-mAb414 at 1:5000, anti-Ran at 1:5000 and anti-α-tubulin at 1:5000 dilutions with overnight incubation at 4°C. HRP-coupled secondary antibody was used at 1:15,000 dilutions for detection of proteins in western blot developed with super signal west Pico chemiluminescent substrate (Pierce, Thermo Fisher Scientific).

### Immunostaining

*In vivo* localization of *d*Elys was revealed by Immunostaining of *Drosophila* embryos. W^1118^ staged syncytial blastoderm embryos were collected on apple juice agar plates for 4 hours at 25°C. Embryos were processed as mentioned by (Harel et al., 1989;Johansen and Johansen, 2004). Briefly, embryos were dechorionated in 50 % sodium hypochlorite until appendages from 80% of embryos disappear. Embryos were washed thoroughly in embryo wash buffer containing 0.2% NaCl and 0.05% Triton X-100 and fixed with 4% formaldehyde in heptane for 45 min at room temperature. Embryos were then devitellinized in 100% methanol for 30 min at room temperature. Embryos were blocked in 5% neutralized goat serum (Jackson Laboratories). Processed embryos were then used for Immunostaining with the anti-*d*Elys antibody (1:1000) and mAb414 (1:500, Bio-legend). Embryos collected from GFP-Nup107 expressing fly line were processed identically. Embryos were mounted in vectashield mounting medium (Vector Laboratories).

For immunostaining of *Drosophila* salivary glands, wandering third instar larva were dissected in cold PBS for isolation of salivary glands. Glands were fixed in freshly prepared 4% formaldehyde for 30 min at room temperature and washed thoroughly with 0.2% PBST (PBS+0.2% Triton X-100). Salivary glands were blocked with 5% neutralized goat serum (Jackson Laboratories) and stained with anti-_d_Elys (1:1000), anti-dNup43 (1:250),mAb414 (1:500, Bio-legend), anti-Ran (1:5000, BD bioscience), anti Lamin Dm0 (1:1000, ADL67.10, DSHB, deposited by Prof. Paul A. Fisher, Stony Brook University, USA (Riemer et al., 1995)), anti-Lamin C (1:2000, gift from Prof. Paul A. Fisher, Stony Brook University, USA (Riemer et al., 1995)), anti-LBR (1:500, gift from Prof. George Krohne, University of Wurzburg, Germany, (Wagner et al., 2004)), anti-Nup98 (1:1000, gift from Dr.Cordula Schulz, University of Georgia, USA (Parrott et al.,2011), anti-TBP (1:200,(Choudhury et al., 2017) for overnight at 4°C. Larva were also probed with anti-Dorsal (1:20, 7A4, DSHB, ((Whalen and Steward, 1993)), anti-Cactus (1:50, 3H12, DSHB,(Whalen and Steward, 1993)). For apoptotic induction, salivary glands were probed with an anti-Dcp-1 antibody (1:100, Cell signaling technology (DeVorkin et al., 2014)). Eye and wing imaginal discs were dissected from third instar larva and processed as described above for probing of apoptotic induction with the Dcp-1 antibody. Secondary antibodies used were anti-rabbit Alexa Fluor 568 (1:800, Thermo Fisher Scientific), anti-mouse Alexa Fluor 488 (1:800, Thermo Fisher Scientific), anti-guinea pig FITC (1:400, Jackson laboratories), anti-mouse Cy5 (1:500, Jackson Laboratories). DNA was stained with DAPI (1:5000, Thermo Fisher Scientific). Salivary glands were mounted in vectashield mounting medium (Vector Laboratories).

*Drosophila* S2 cells were immunostained by immobilizing on 0.25 mg/ml concanavalin A coated coverslips for 2 hours at 25°C. MG-132 arrested cells were fixed by 100% methanol at −20°C, blocked with 5% neutralized goat serum (Jackson laboratories) and immunostained anti-*d*Elys (1:1000), anti-α-tubulin (1:1000, DSHB), anti-CID (1:1000, gift from Prof. Steve Henikoff, FHCRC,USA, (Vermaak et al., 2002)). All the steps of immunostaining of S2 cells were followed as per (Buster et al., 2010).

All the samples were imaged on Carl Zeiss LSM 780 up-right confocal microscope equipped with 63X/1.4 N.A. oil immersion lens. Super-resolution microscopy of anti-CID antibody labeled, EYFP-*d*Elys transfected cells were done with Carl Zeiss LSM 800 Airyscan inverted microscope. Images were processed for super-resolution in the *inbuilt* mathematical algorithm of Zen.2 software of LSM 800 Airyscan. Images were processed with ImageJ (NIH) or Fiji software and Adobe Photoshop CS6 (Adobe Corporation). Nuclei sizes were measured by using Fiji software and graph were plotted with GraphPad software (Prism).

### Acridine Orange staining for apoptosis

To examine the level of apoptosis in control as well as *d*Elys knockdown salivary glands, dissected glands were incubated with 2 μg/ml acridine orange (Sigma) for 5 min at room temperature. Glands were imaged immediately without fixation with Carl Zeiss LSM 780 up-right confocal microscope (Denton et al., 2008; Sarkissian et al., 2014). Fluorescence intensities of the nucleus were calculated using Fiji software. ~150 nuclei from three independent experiments were counted and the graph was plotted in GraphPad software (Prism).

### Bright field microscopy

For imaging of eye and wing phenotype of *d*Elys knockdown flies, three days old flies were anesthetized with diethyl ether (Merck) and immobilized on sticky gum for proper orientation of *Drosophila* eye. At least 10 flies from each knockdown experiment were imaged from three independent experiments for each genotype. *Drosophila* eyes were imaged using Leica fluorescent stereo-microscope M205 FA using a 123X magnification of particular equipment. For wing imagining in case of wing phenotypes with a *d*Elys knockdown, three-day-old fly wings from each genotype were dissected and placed on a glass slide under a coverslip and imaged directly under Leica upright light microscope DM2500 with 10X magnification (Velentzas et al., 2013). At least 15 pairs of wings from each genotype were examined from three independent crosses.

### Scanning electron microscopy (SEM)

*Drosophila* eyes from eye-specific knockdown of *d*Elys were imaged using a scanning electron microscope for detailed analysis of eye structure perturbations. Three days old flies from each RNAi cross was processed as per Wolff, 2011). Briefly, flies were anesthetized and fixed with 2.5% glutaraldehyde in phosphate buffered saline for 2 hours at 4°C. Flies were washed thoroughly twice with PBS+ 4% sucrose. Flies were dehydrated through a graded ethanol series and were subjected to critical point drying. Samples were mounted on aluminum stubs with carbon conductive tape. Flies were coated with gold particles in a sputter coating apparatus. Samples were imaged using Carl Zeiss Gemini II FESEM microscope. At least 10 flies from each genotype were imaged from three independent experiments.

### *In vitro* DNA binding experiment

To analyze DNA binding activity of AT-hook motifs of *d*Elys, we purified fragment spanning all three AT-hook DNA binding motif by cloning C-terminal 104 amino acid (1858-1962) from clone LD14710 into pET28a(+) vector. (His)_6_ tagged AT-hook motif fragment was purified by expressing in *E.coli* BL21 (DE3) cells. Protein was induced by 200 μM IPTG (Sigma) overnight at 18°C. Cells were lysed in lysis buffer containing 50 mM Tris-HCl pH8.0, 150 mM NaCl, 20 mM imidazole, 1% Triton X-100, and 50 μg/ml lysozyme on ice. Protein was purified over Ni-NTA beads by eluting in 300 mM imidazole in 50 mM Tris-HCl pH 8.0, 150 mM NaCl, 1 mM MgCl_2_ and 0.5 mM EDTA. Protein was dialyzed overnight against dialysis buffer, 50 mM Tris-HCl pH 8.0, 150 mM NaCl, 1 mM MgCl_2_ and 0.5 mM EDTA.

For site-directed mutagenesis of arginine residues, we used Q5 site-directed mutagenesis kit (NEB Bio-labs) with primers coding mutated nucleotides. Primers used for mutagenesis were mentioned in Table S2. The clone described above was used as a template for site-directed mutagenesis. Mutations were confirmed using DNA sequencing (IISER-Bhopal sequencing facility). Clones were transformed into *E.coli* BL21 (DE3) cells and proteins were purified as mentioned earlier.

*In vitro* DNA binding experiment was performed by incubating template which contains 50% AT richness and 90% AT richness with purified wild-type and mutant proteins. A known cytosolic protein which does not binds to DNA as a negative control. DNA binding experiment was performed in binding buffer containing 100 mM Tris, 500 mM KCl, 10 mM DTT pH 7.5 for 30 min at room temperature and analyzed on to 0.8% agarose gel and detected by UV transilluminator (UVP). Purified DNA used for the binding experiment is 5 nM and total purified protein was used at 0.5 μM concentrations. EMSA was performed by using Lightshift Chemiluminescent EMSA Kit (Pierce, Thermo Scientific) by using *Sdic* DNA oligos as described by (Metcalf and Wassarman, 2006).

### Quantitative-PCR

Total RNA was isolated from Control and *d*Elys knockdown third instar larva head complex using total tissue RNA isolation kit (Favorgen Biotech). One μg of total RNA was used to synthesize cDNA using iScript cDNA synthesis (Bio-Rad). cDNA was diluted 5 times and 1 μl of cDNA from each genotype were used as a template for semi-quantitative PCR using *d*Elys specific primers and RpL49 as a control. Real-time PCR on the same cDNA was done in Roche Lightcycler 480 at standard cycling conditions and probed with SYBR-green (Bio-Rad) in real time using *d*Elys and Actin RT primers. Quantification of each reaction was done by calculating ΔC^T^ values. ΔC^T^ was normalized against C^T^ values of actin. The graph was plotted as fold change in *d*Elys expression using GraphPad software (Prism). Primers used in quantitative PCR are mentioned in table S2.

## Acknowledgment

We thank Bloomington Drosophila Stock Centre (BDSC) and Vienna *Drosophila* Resource Centre (VDRC) for fly lines used in this study, *Drosophila* Genomics Resource Centre (DGRC) for a genomic clone of CG14215 and Developmental Studies Hybridoma Bank (DSHB), the University of Iowa for monoclonal antibodies. We thank Prof. Paul A. Fisher, Stony Brook University for Lamin C antibodies, Prof. George Krohne, the University of Wurzburg for LBR antibody, Dr. Cordula Schulz, the University of Georgia for the Nup98 antibody, Prof. Steve Henikoff, FHCRC, USA for CID antibody and Dr. R.S.Tomar, IISER-Bhopal for TBP antibody. We thank IISER-Bhopal Central Instrumentation Facility for DNA sequencing, SEM and Confocal Microscopes.

## Author Contributions

Conceptualization: RKM and SJM. Methodology: SJM, RKM and VK; Formal Analysis: SJM, RKM and VK; Investigation: SJM; Resources: RKM and VK; Writing – original draft: SJM and RKM; Writing - review & editing: SJM, RKM and VK; Visualization: SJM; Supervision: RKM and VK; Project administration: RKM; Funding acquisition: RKM

---

**Table S1:** Fly line used in this study.

| Line No. | Genotype | Origin |
| --- | --- | --- |
| BL-35514 | w[*];P{w[+mC]=GFP-Nup107.K}9.1 | (Katsani et al., 2008) |
| dElys<sup>KK</sup><br>(v103547) | P{KK100932}VIE-260B | VDRC |
| dElys<sup>TRiP</sup> | Y<sup>1</sup>V<sup>1</sup>;p{pVALIUM10_dElys-1}/Cyo;attp40 | This Study (TRiP based) |
| EYFP-dElys | W[*],p{pTVW-dElys} | This Study |
| BL-53269 | y1 sc* v1; P{TRiP.HMC02426}attP2 | (Ni et al., 2011) |
| BL-4776 | w1118; P{UAS-GFP.nls}8 | BDSC |
| BL-1691 | y[1] w[67c23]; P{w[+mC]=Ubi-GFP.nls}ID-2; P{Ubi-GFP.nls}ID-3 | BDSC |
|  | dElys<sup>KK</sup>; Ubi-GFP-NLS | This Study |
| BL-25374 | y1 w*; P{Act5C-GAL4-w}E1/CyO | BDSC |
| BL-5534 | w*; P{GAL4-ey.H}3-8 | BDSC |
| BL-25754 | P{UAS-Dcr-2.D}1, w1118; P{GawB}nubbin-AC-62 | BDSC |
| BL-1561 | w*; P{GAL4-arm.S}4a P{GAL4-arm.S}4b/TM3, Sb1 Ser1 | BDSC |
| BL-7062 | w[*]; P{w[+mC]=matalpha4-GAL-VP16}V2H | BDSC |
| BL-6657 | w[*]; P{w[+mC]=UAS-DIAP1.H}3 | BDSC |
|  | dElys<sup>KK</sup>; UAS-DIAP1 | This Study |
| v105491<br>(Dorsal RNAi) | P{KK107820}VIE-260B | VDRC |
| | $dElys^{KK}; Dorsal^{RNAi}$ | This Study |
| v21937<br>(Nup160 RNAi) | w[1118]; P{GD11413}v21937 | VDRC |
| v110194<br>(Nup133 RNAi) | P{KK102392}VIE-260B | VDRC |
| V22407<br>(Nup107 RNAi) | w[1118]; P{GD12024}v22407 | VDRC |
| BL-32837<br>(Nup153 RNAi) | y[1] sc[*] v[1]; P{y[+t7.7]<br>v[+t1.8]=TRiP.HMS00527}attP2 | BDSC |
| BL-47326<br>(fkh-GAL4) | w[1118]; P{y[+t7.7] w[+mC]=GMR16E04-GAL4}attP2 | BDSC |
| BL-6979<br>(C147-GAL4) | w[1118]; P{w[+mW.hs]=GawB}C147 | BDSC |
|  | w[1118];C147-GAL4;fkh-GAL4 | This study |

**Table S2:**
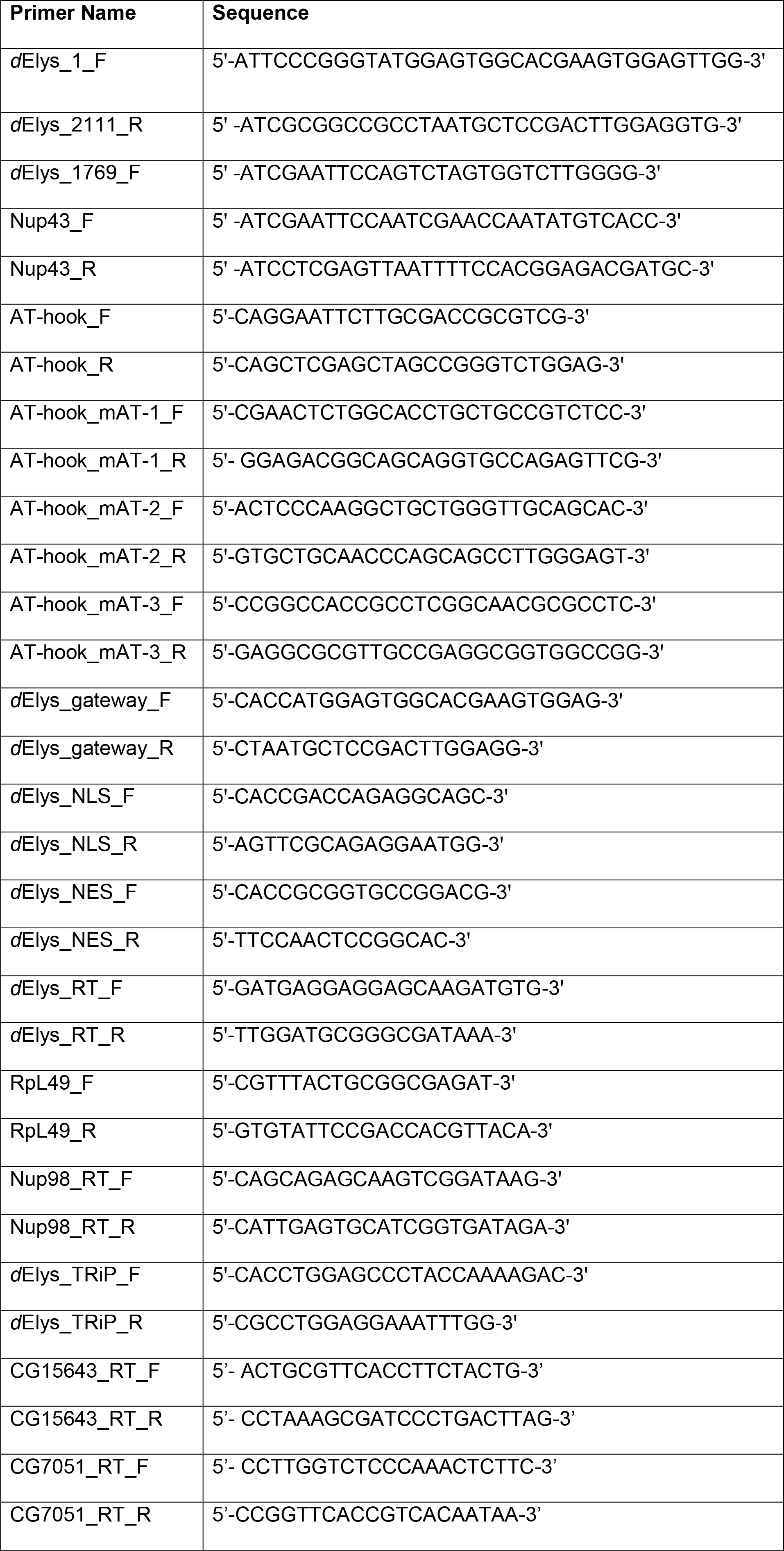
Primer used in this study.

**Figure S1:**
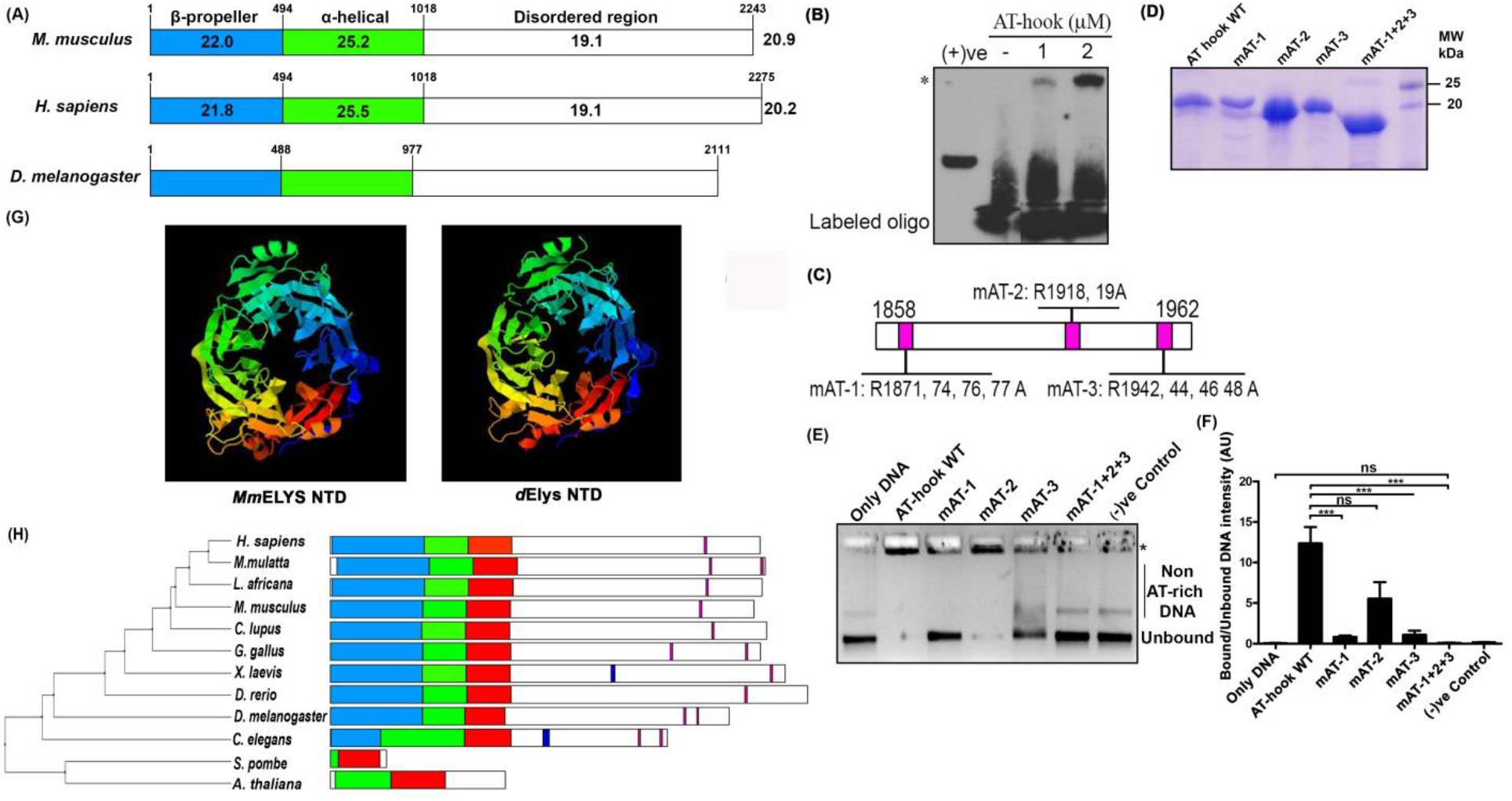
*d*Elys *in silico* characterization and EMSA. (A) Graphical representation of sequence identity of *d*Elys with that of Human and mouse ELYS. *d*Elys is 20.2% and 20.9% identical with human and mouse ELYS respectively. Domain wise identity is mentioned above each domain highlighted. Conserved secondary structures between *Drosophila*, mouse and human ELYS are represented. *d*Elys contains β-sheets which form the β-propeller structure as identical as a mouse and human ELYS N-terminal domain. β-propeller N-terminal domain, α-helical central domain, and disordered C-terminal regions are conserved in *d*Elys as in higher orthologues. (B) Assessment of DNA binding activity of *d*Elys wild-type AT-hook motifs performed by EMSA. The asterisk represents the super-shifted DNA-AT-hook complex. (C) Graphical representation of the position of each AT-hook motif of *d*Elys in fragment cloned for protein purification. Conserved arginine residues were mutated to alanine and the position of each arginine residues are indicated. (D) SDS-PAGE analysis of purified wild type and mutant AT-hook proteins used for DNA binding experiment. Molecular weight markers are indicated next to the gel. (E) Non-AT rich DNA binding ability of predicted AT-hooks is tested in EMSA experiment. Variant of protein used in the reaction is indicated above each lane. Asterisk indicates bound DNA with decreased mobility. (F) Quantification of DNA binding ability of each AT-hook fragment used. Quantification is derived from at least three independent experiments. *** represents p<0.0001, ns is non-significant. Error bar represents a Standard error of the mean (SEM). Statistical analysis is derived from one-way ANOVA followed by post-hoc Tukey’s multiple comparison tests. (G) N-terminal domain of *d*Elys showing identical seven-bladed β-propeller structure as known for the N-terminal domain of Mouse ELYS (mention about the prediction of the structure). (H) Phylogenetic analysis of ELYS orthologues from yeast, invertebrate, vertebrates, and plant. Domain structure of ELYS represented as β-sheet containing N-terminal domain (blue), a-helical central domain (green), conserved ELYS domain (red), coiled-coil domain (dark blue, ensure that it is coiled-coil region) and AT-hook DNA binding motifs (purple).

**Figure S2:**
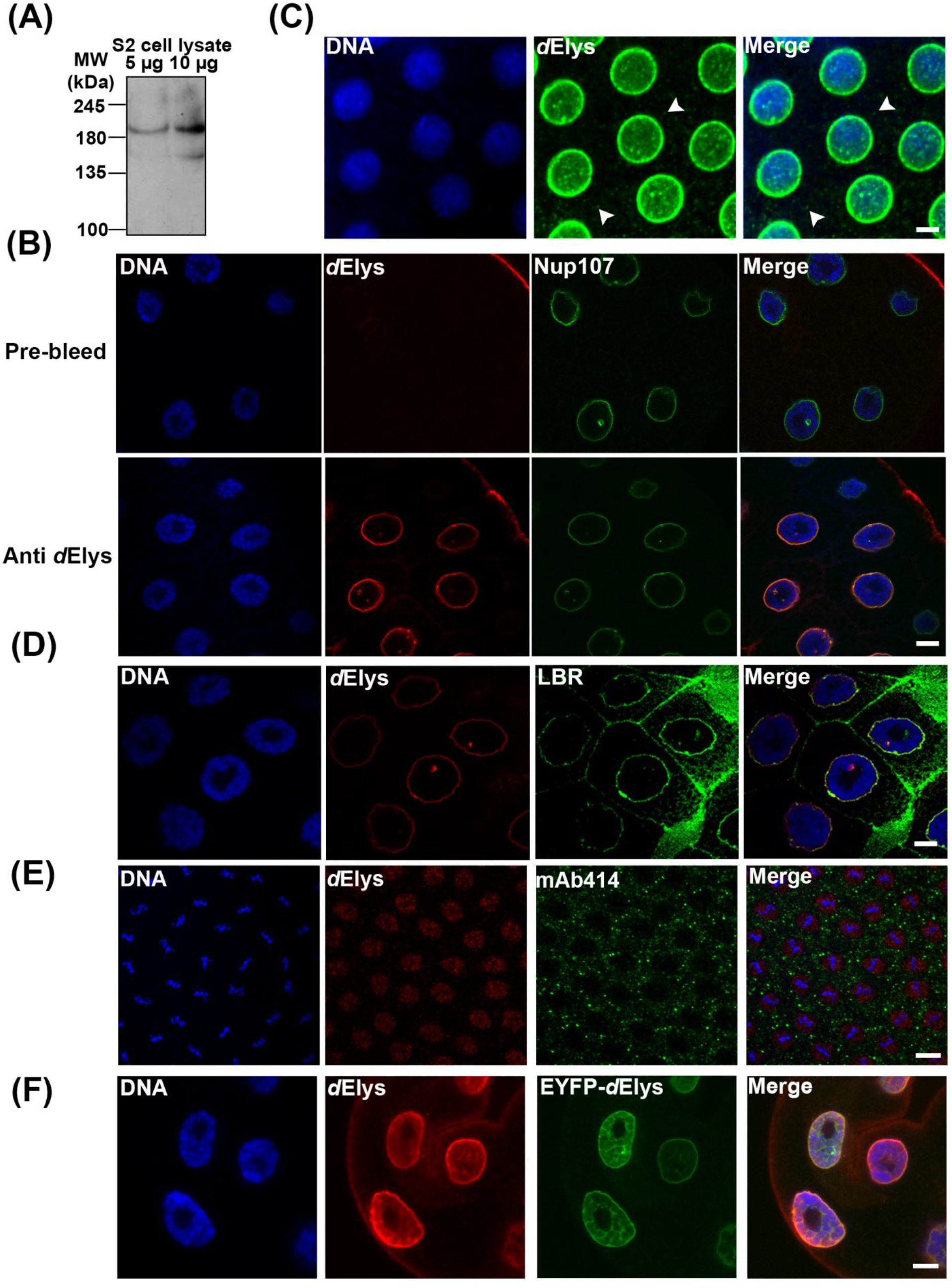
*d*Elys antibody characterization and its localization *in vivo*. (A) Anti-*d*Elys antibody picks a band of ~235 kDa in *Drosophila* S2 cell lysate. It also detects degradation bands of *d*Elys. (B) The pre-bleed serum and anti-*d*Elys serum (red) used on salivary gland from GFP-Nup107 (green) expressing larva. DNA is stained with DAPI. (Scale bar: 10 μm) (C) Presence of *d*Elys in cytoplasmic structure, annulate lamellae (AL) visualizedible under saturating condition of imaging. DNA is stained with DAPI. (Scale bar: 5 μm) (D) Localization of *d*Elys (red) and Lamin B receptor (LBR) (green) at nuclear rim in *Drosophila* salivary glands. DNA is stained with DAPI. (Scale bar: 5 μm) (E) Early *Drosophila* mitotic syncytial embryo stained with anti-*d*Elys (red) antibody mAb414 (green) marks mitotic cytoplasm with little presence around mitotic chromosome. DNA is stained with DAPI. (Scale bar: 5 μm) (F) EYFP tagged *d*Elys (green) expressed in *Drosophila* salivary gland nuclei DNA is stained with DAPI. (Scale bar: 5 μm)

**Figure S3:**
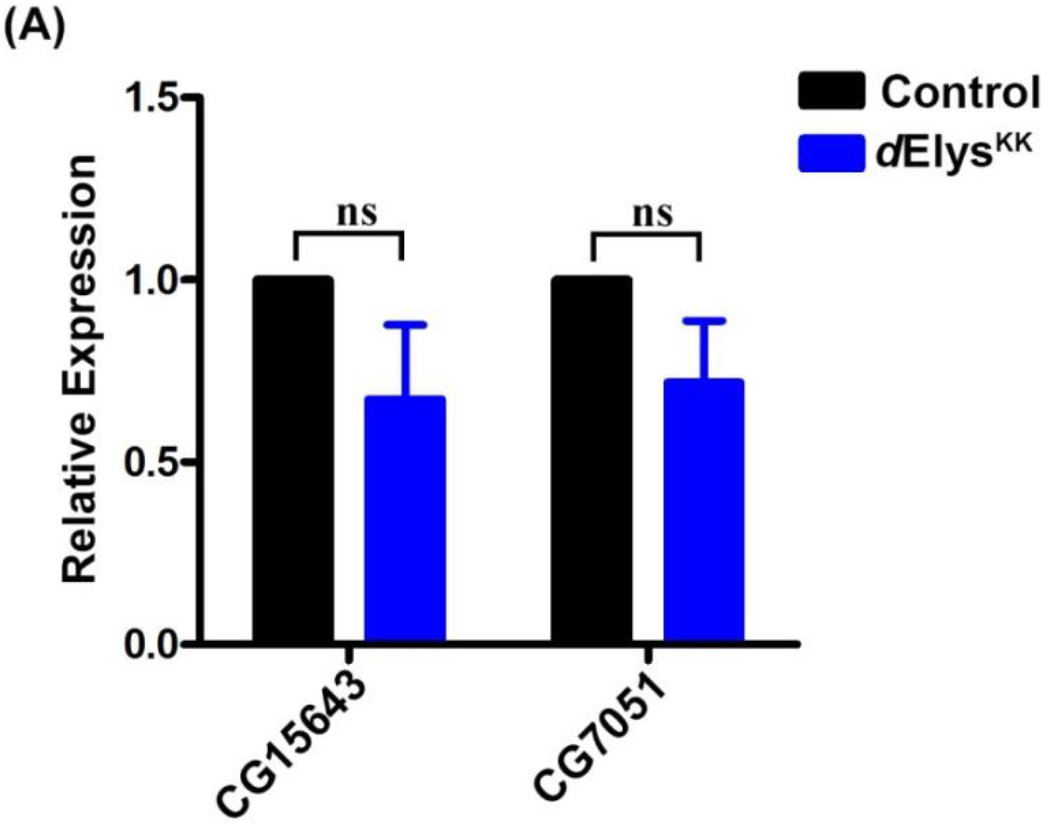
No change in expression of predicted off-targets of *d*Elys^KK^ RNAi line. (A) Quantitative PCR showing the expression level of predicted off-targets of *d*Elys^KK^ RNAi line from VDRC. Data is represented from at least three independent experiments. Statistical significance derived from student’s t-test. Error bar represents standard deviation. ns is non-significant.

**Figure S4:**
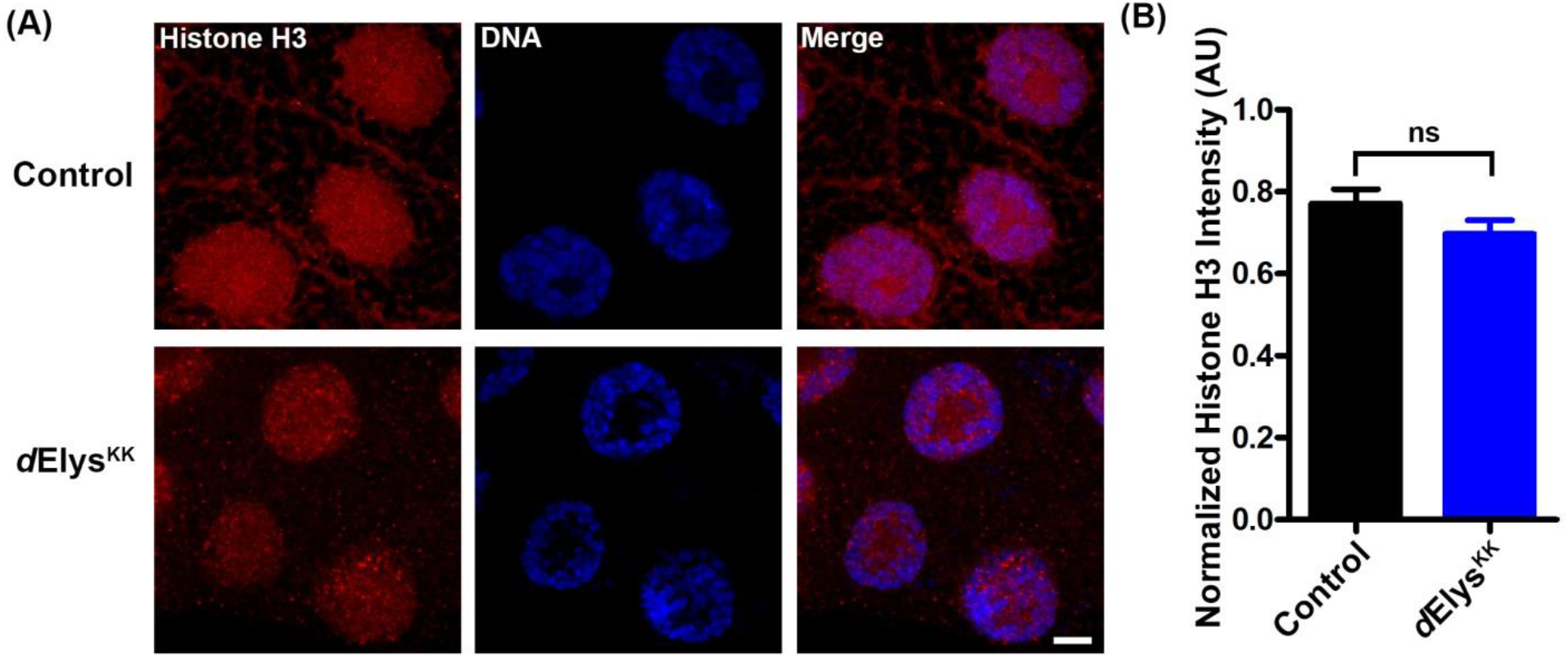
*d*Elys depletion does not affect Histone-H3 recruitment to nucleosome. (A) Representative images showing histone H3 (red) staining in control and *d*Elys depleted (*Act*5C-GAL4 driven) salivary gland nuclei. DNA is stained with DAPI. (Scale bar: 5 μm) (B) Quantification of Histone H3 intensity compared in control and *d*Elys RNAi. Data is represented from at least three independent experiments. Statistical significance derived from student’s t-test. Error bar represents SEM. ns is non-significant.

**Figure S5:**
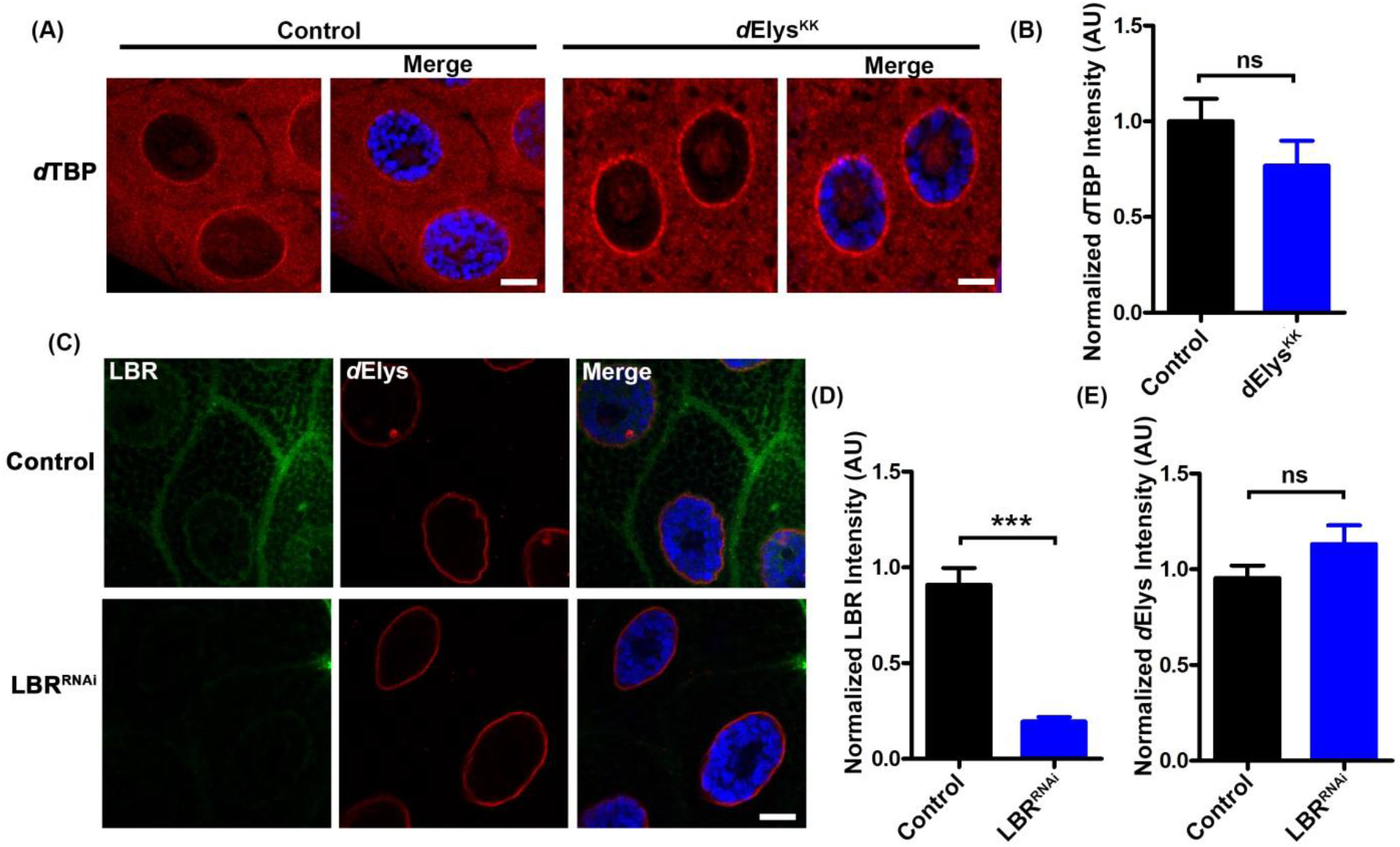
Nuclear membrane is not perturbed by *d*Elys depletion and *d*Elys is not affected by LBR depletion. (A) Staining of nuclear membrane-associated protein ďTBP in control and *d*Elys depleted (*Act*5C-GAL4 driven) salivary gland nuclei. (Scale bar: 5 μm) (B) Quantitation of nuclear rim localization of ďTBP in control and *d*Elys RNAi. Intensities were normalized against the intensity of DAPI. Data is represented from at least three independent experiments. Statistical significance derived from student’s t-test. Error bar represents SEM. ns is non-significant. (C) Salivary gland nuclei showing nuclear rim localization of *d*Elys in LBR RNAi (*Act*5C-GAL4 driven). LBR is in green, *d*Elys in red. DNA is stained with DAPI. (Scale bar: 5 μm) (D, E) Quantitation of nuclear rim intensity of each indicated molecules in control and LBR RNAi. The intensity of each molecule is normalized against the intensity of DAPI. Data is represented from at least three independent experiments. Statistical significance derived from student’s t-test. Error bar represents SEM. *** represents p<0.0001, ns is non-significant.

**Figure S6:**
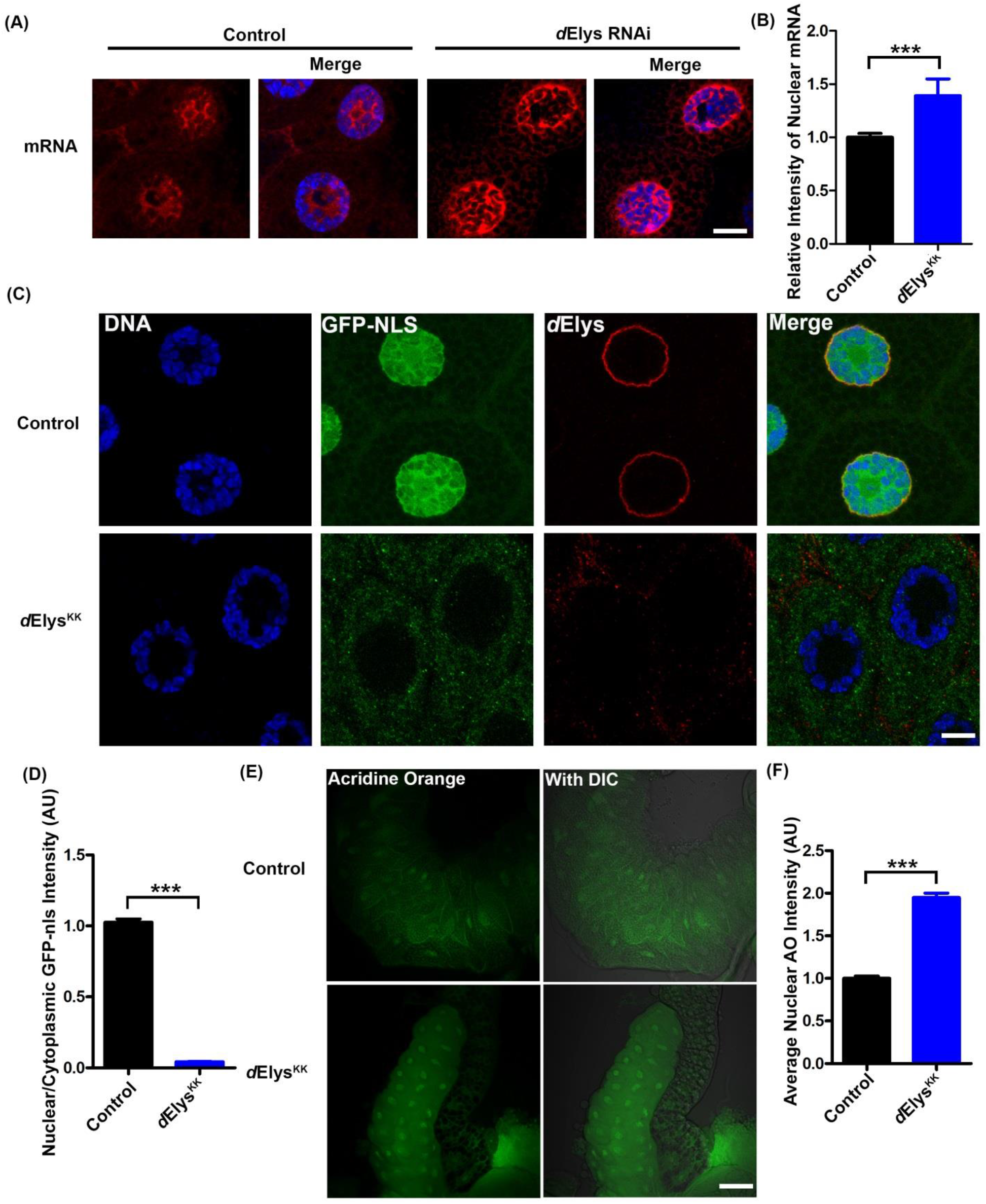
*d*Elys depletion affects nucleocytoplasmic transport and induces apoptosis. (A) Nuclear import of GFP-NLS (green) in control and *d*Elys silenced (*Act*5C-GAL4 driven) salivary gland nuclei. *d*Elys is stained in red. DNA is stained with DAPI. (Scale bar: 5 μm) (B) Acridine orange (green) staining of salivary gland isolated from third instar larva in control and *d*Elys knockdown (*Act*5C-GAL4 driven). DIC is for reference to the presence of salivary gland. (Scale bar: 50 μm) (C) Quantitative analysis of nuclear acridine orange intensities in control and *d*Elys knockdown as seen in (B). Data is represented from at least three independent experiments. Statistical significance derived from student’s t-test. Error bar represents standard error of the mean. ***p<0.0001 (D) Quantitative analysis of nuclear to cytoplasmic GFP-NLS intensities as seen in (A). Data is represented from at least three independent experiments. Statistical significance derived from student’s t-test. Error bar represents standard error of the mean. ***p<0.0001

**Figure S7:**
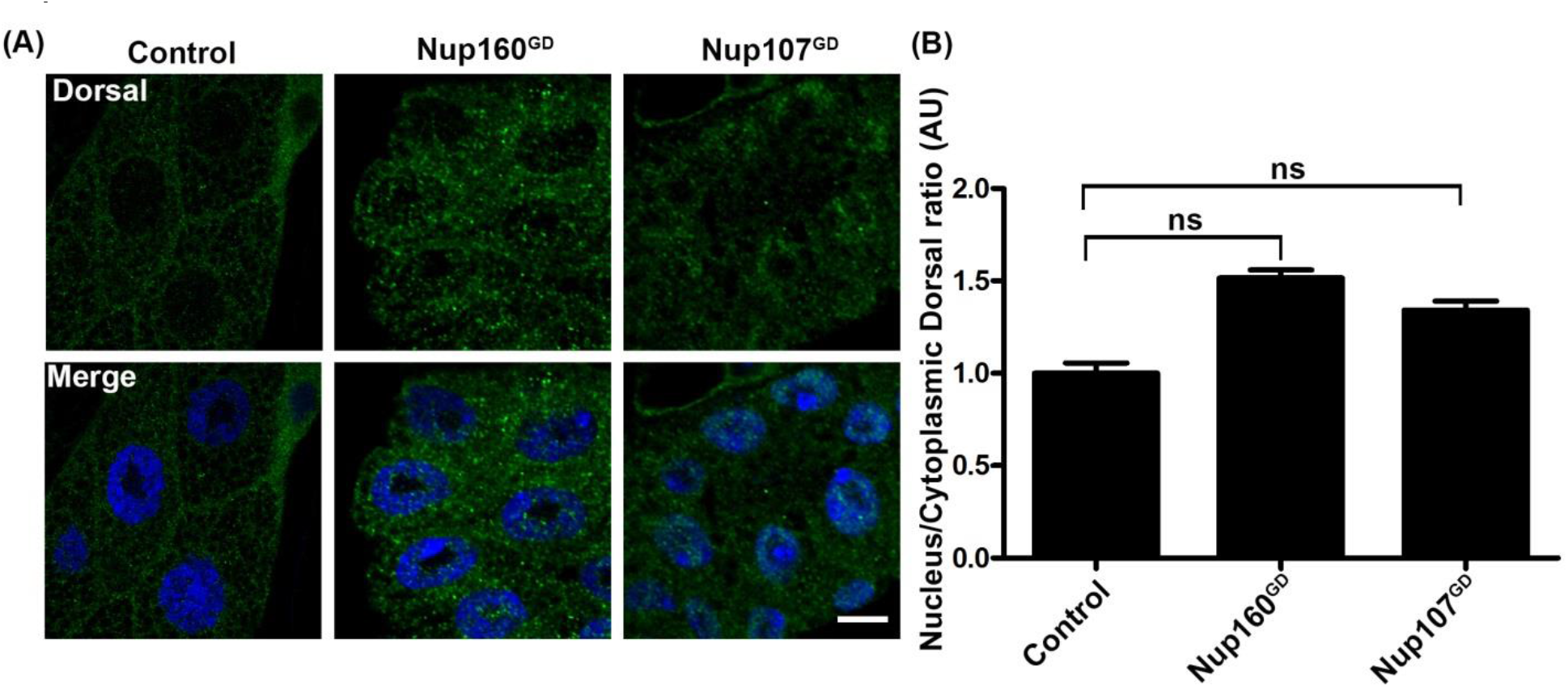
Dorsal nuclear localization is not evident in Nup160 and Nup107 depletion. (A) Nuclear Dorsal presence assessed in nucleoporin knockdown salivary gland nuclei. DNA is stained with DAPI. (Scale bar: 5 μm) (B) Quantitation of nuclear/cytoplasmic intensity ratio of dorsal in nucleoporin knockdown nuclei. Data is represented from at least three independent experiments. Statistical significance derived from one way ANOVA followed by Tukey’s post-hoc test. Error bar is SEM. ns is non-significant.

## References

Aitchison, J. D., Blobel, G. and Rout, M. P. (1995). Nup120p: a yeast nucleoporin required for NPC distribution and mRNA transport. J Cell Biol 131, 1659–75.

Andlauer, T. F., Scholz-Kornehl, S., Tian, R., Kirchner, M., Babikir, H. A., Depner, H., Loll, B., Quentin, C., Gupta, V. K., Holt, M. G. et al. (2014). Drep-2 is a novel synaptic protein important for learning and memory. Elife 3.

Aravind, L. and Landsman, D. (1998). AT-hook motifs identified in a wide variety of DNA-binding proteins. Nucleic Acids Res 26, 4413–21.

Barkett, M. and Gilmore, T. D. (1999). Control of apoptosis by Rel/NF-kappaB transcription factors. Oncogene 18, 6910–24.

Belvin, M. P., Jin, Y. and Anderson, K. V. (1995). Cactus protein degradation mediates Drosophila dorsal-ventral signaling. Genes Dev 9, 783–93.

Bilokapic, S. and Schwartz, T. U. (2013). Structural and functional studies of the 252 kDa nucleoporin ELYS reveal distinct roles for its three tethered domains. Structure 21, 572–80.

Buster, D. W., Nye, J., Klebba, J. E. and Rogers, G. C. (2010). Preparation of Drosophila S2 cells for light microscopy. J Vis Exp.

Campbell, K., Lebreton, G., Franch-Marro, X. and Casanova, J. (2018). Differential roles of the Drosophila EMT-inducing transcription factors Snail and Serpent in driving primary tumour growth. PLoS Genet 14, e1007167.

Cerveny, K. L., Cavodeassi, F., Turner, K. J., de Jong-Curtain, T. A., Heath, J. K. and Wilson, S. W. (2010). The zebrafish flotte lotte mutant reveals that the local retinal environment promotes the differentiation of proliferating precursors emerging from their stem cell niche. Development 137, 2107–15.

Choudhury, S. D., Vs, A., Mushtaq, Z. and Kumar, V. (2017). Altered translational repression of an RNA-binding protein, Elav by AOA2-causative Senataxin mutation. Synapse 71.

Clever, M., Funakoshi, T., Mimura, Y., Takagi, M. and Imamoto, N. (2012). The nucleoporin ELYS/Mel28 regulates nuclear envelope subdomain formation in HeLa cells. Nucleus 3, 187–99.

Davuluri, G., Gong, W., Yusuff, S., Lorent, K., Muthumani, M., Dolan, A. C. and Pack, M. (2008). Mutation of the zebrafish nucleoporin elys sensitizes tissue progenitors to replication stress. PLoS Genet 4, e1000240.

de Jong-Curtain, T. A., Parslow, A. C., Trotter, A. J., Hall, N. E., Verkade, H., Tabone, T., Christie, E. L., Crowhurst, M. O., Layton, J. E., Shepherd, I. T. et al. (2009). Abnormal nuclear pore formation triggers apoptosis in the intestinal epithelium of elys-deficient zebrafish. Gastroenterology 136, 902–11.

DeLotto, R., DeLotto, Y., Steward, R. and Lippincott-Schwartz, J. (2007). Nucleocytoplasmic shuttling mediates the dynamic maintenance of nuclear Dorsal levels during Drosophila embryogenesis. Development 134, 4233–41.

Denton, D., Mills, K. and Kumar, S. (2008). Methods and protocols for studying cell death in Drosophila. Methods Enzymol 446, 17–37.

DeVorkin, L., Go, N. E., Hou, Y. C., Moradian, A., Morin, G. B. and Gorski, S. M. (2014). The Drosophila effector caspase Dcp-1 regulates mitochondrial dynamics and autophagic flux via SesB. J Cell Biol 205, 477–92.

Emtage, J. L., Bucci, M., Watkins, J. L. and Wente, S. R. (1997). Defining the essential functional regions of the nucleoporin Nup145p. J Cell Sci 110 (Pt 7), 911–25.

Espin-Palazon, R. and Traver, D. (2016). The NF-kappaB family: Key players during embryonic development and HSC emergence. Exp Hematol 44, 519–27.

Fabre, E. and Hurt, E. (1997). Yeast genetics to dissect the nuclear pore complex and nucleocytoplasmic trafficking. Annu Rev Genet 31, 277–313.

Fernandez, A. G. and Piano, F. (2006). MEL-28 is downstream of the Ran cycle and is required for nuclear-envelope function and chromatin maintenance. Curr Biol 16, 1757–63.

Flockhart, I. T., Booker, M., Hu, Y., McElvany, B., Gilly, Q., Mathey-Prevot, B., Perrimon, N. and Mohr, S. E. (2012). FlyRNAi.org--the database of the Drosophila RNAi screening center: 2012 update. Nucleic Aciďs Res 40, D715–9.

Franz, C., Walczak, R., Yavuz, S., Santarella, R., Gentzel, M., Askjaer, P., Galy, V., Hetzer, M., Mattaj, I. W. and Antonin, W. (2007). MEL-28/ELYS is required for the recruitment of nucleoporins to chromatin and postmitotic nuclear pore complex assembly.EMBO Rep 8, 165–72.

Galy, V., Askjaer, P., Franz, C., Lopez-Iglesias, C. and Mattaj, I. W. (2006). MEL-28, a novel nuclear-envelope and kinetochore protein essential for zygotic nuclear-envelope assembly in C. elegans. Curr Biol 16, 1748–56.

Gao, N., Davuluri, G., Gong, W., Seiler, C., Lorent, K., Furth, E. E., Kaestner, K. H. and Pack, M. (2011). The nuclear pore complex protein Elys is required for genome stability in mouse intestinal epithelial progenitor cells. Gastroenterology 140, 1547–55 e10.

Genenncher, B., Wirthmueller, L., Roth, C., Klenke, M., Ma, L., Sharon, A. and Wiermer, M. (2016). Nucleoporin-Regulated MAP Kinase Signaling in Immunity to a Necrotrophic Fungal Pathogen. Plant Physiol 172, 1293–1305.

Gillespie, P. J., Khoudoli, G. A., Stewart, G., Swedlow, J. R. and Blow, J. J. (2007). ELYS/MEL-28 chromatin association coordinates nuclear pore complex assembly and replication licensing. Curr Biol 17, 1657–62.

Gillespie, S. K. and Wasserman, S. A. (1994). Dorsal, a Drosophila Rel-like protein, is phosphorylated upon activation of the transmembrane protein Toll. Mol Cell Biol 14, 3559–68.

Gomez-Saldivar, G., Fernandez, A., Hirano, Y., Mauro, M., Lai, A., Ayuso, C., Haraguchi, T., Hiraoka, Y., Piano, F. and Askjaer, P. (2016). Identification of Conserved MEL-28/ELYS Domains with Essential Roles in Nuclear Assembly and Chromosome Segregation. PLoS Genet 12, e1006131.

Govind, S. (1999). Control of development and immunity by rel transcription factors in Drosophila. Oncogene 18, 6875–87.

Gullaud, M., Delanoue, R. and Silber, J. (2003). A Drosophila model to study the functions of TWIST orthologs in apoptosis and proliferation. Cell Death Differ 10, 641–51.

Harel, A., Zlotkin, E., Nainudel-Epszteyn, S., Feinstein, N., Fisher, P. A. and Gruenbaum, Y. (1989). Persistence of major nuclear envelope antigens in an envelope-like structure during mitosis in Drosophila melanogaster embryos. J Cell Sci 94 (Pt 3), 463–70.

Hattersley, N., Cheerambathur, D., Moyle, M., Stefanutti, M., Richardson, A., Lee, K. Y., Dumont, J., Oegema, K. and Desai, A. (2016). A Nucleoporin Docks Protein Phosphatase 1 to Direct Meiotic Chromosome Segregation and Nuclear Assembly. Dev Cell 38, 463–77.

Hetzer, M. W. (2010). The nuclear envelope. Colď Spring Harb Perspect Biol 2, a000539.

Hoelz, A., Debler, E. W. and Blobel, G. (2011). The structure of the nuclear pore complex. Annu Rev Biochem 80, 613–43.

Ilyin, A. A., Ryazansky, S. S., Doronin, S. A., Olenkina, O. M., Mikhaleva, E. A., Yakushev, E. Y., Abramov, Y. A., Belyakin, S. N., Ivankin, A. V., Pindyurin, A. V. et al.(2017). Piwi interacts with chromatin at nuclear pores and promiscuously binds nuclear transcripts in Drosophila ovarian somatic cells. Nucleic Aciďs Res 45, 7666–7680.

Johansen, K. M. and Johansen, J. (2004). Studying nuclear organization in embryos using antibody tools. Methoďs Mol Biol 247, 215–34.

Jones, D. T. (1999). Protein secondary structure prediction based on position-specific scoring matrices. J Mol Biol 292, 195–202.

Kaltschmidt, B., Kaltschmidt, C., Hofmann, T. G., Hehner, S. P., Droge, W. and Schmitz, M. L. (2000). The pro-or anti-apoptotic function of NF-kappaB is determined by the nature of the apoptotic stimulus. Eur J Biochem 267, 3828–35.

Katsani, K. R., Karess, R. E., Dostatni, N. and Doye, V. (2008). In vivo dynamics of Drosophila nuclear envelope components. Mol Biol Cell 19, 3652–66.

Kelley, L. A., Mezulis, S., Yates, C. M., Wass, M. N. and Sternberg, M. J. (2015). The Phyre2 web portal for protein modeling, prediction and analysis. Nat Protoc 10, 845–58.

Kimura, N., Takizawa, M., Okita, K., Natori, O., Igarashi, K., Ueno, M., Nakashima, K., Nobuhisa, I. and Taga, T. (2002). Identification of a novel transcription factor, ELYS, expressed predominantly in mouse foetal haematopoietic tissues. Genes Cells 7, 435–46.

Kiseleva, E., Rutherford, S., Cotter, L. M., Allen, T. D. and Goldberg, M. W. (2001). Steps of nuclear pore complex disassembly and reassembly during mitosis in early Drosophila embryos. J Cell Sci 114, 3607–18.

Lim, H. Y. and Tomlinson, A. (2006). Organization of the peripheral fly eye: the roles of Snail family transcription factors in peripheral retinal apoptosis. Development 133, 3529–37.

Liu, W., Xie, Y., Ma, J., Luo, X., Nie, P., Zuo, Z., Lahrmann, U., Zhao, Q., Zheng, Y., Zhao, Y. et al. (2015). IBS: an illustrator for the presentation and visualization of biological sequences. Bioinformatics 31, 3359–61.

McCall, K. and Steller, H. (1998). Requirement for DCP-1 caspase during Drosophila oogenesis. Science 279, 230–4.

Metcalf, C. E. and Wassarman, D. A. (2006). DNA binding properties of TAF1 isoforms with two AT-hooks. J Biol Chem 281, 30015–23.

Mimura, Y., Takagi, M., Clever, M. and Imamoto, N. (2016). ELYS regulates the localization of LBR by modulating its phosphorylation state. J Cell Sci 129, 4200–4212.

Mishra, R. K., Chakraborty, P., Arnaoutov, A., Fontoura, B. M. and Dasso, M. (2010). The Nup107-160 complex and gamma-TuRC regulate microtubule polymerization at kinetochores. Nat Cell Biol 12, 164–9.

Neisch, A. L., Speck, O., Stronach, B. and Fehon, R. G. (2010). Rho1 regulates apoptosis via activation of the JNK signaling pathway at the plasma membrane. J Cell Biol 189, 311–23.

Ni, J. Q., Zhou, R., Czech, B., Liu, L. P., Holderbaum, L., Yang-Zhou, D., Shim, H. S., Tao, R., Handler, D., Karpowicz, P. et al. (2011). A genome-scale shRNA resource for transgenic RNAi in Drosophila. Nat Methods 8, 405–7.

Notredame, C., Higgins, D. G. and Heringa, J. (2000). T-Coffee: A novel method for fast and accurate multiple sequence alignment. J Mol Biol 302, 205–17.

Okita, K., Kiyonari, H., Nobuhisa, I., Kimura, N., Aizawa, S. and Taga, T. (2004). Targeted disruption of the mouse ELYS gene results in embryonic death at peri-implantation development. Genes Cells 9, 1083–91.

Okita, K., Nobuhisa, I., Takizawa, M., Ueno, M., Kimura, N. and Taga, T. (2003). Genomic organization and characterization of the mouse ELYS gene. Biochem Biophys Res Commun 305, 327–32.

Ori, A., Banterle, N., Iskar, M., Andres-Pons, A., Escher, C., Khanh Bui, H., Sparks, L., Solis-Mezarino, V., Rinner, O., Bork, P. et al. (2013). Cell type-specific nuclear pores: a case in point for context-dependent stoichiometry of molecular machines. Mol Syst Biol 9, 648.

Orjalo, A. V., Arnaoutov, A., Shen, Z., Boyarchuk, Y., Zeitlin, S. G., Fontoura, B., Briggs, S., Dasso, M. and Forbes, D. J. (2006). The Nup107-160 nucleoporin complex is required for correct bipolar spindle assembly. Mol Biol Cell 17, 3806–18.

Parrott, B. B., Chiang, Y., Hudson, A., Sarkar, A., Guichet, A. and Schulz, C. (2011). Nucleoporin98-96 function is required for transit amplification divisions in the germ line of Drosophila melanogaster. PLoS One 6, e25087.

Pascual-Garcia, P., Debo, B., Aleman, J. R., Talamas, J. A., Lan, Y., Nguyen, N. H., Won, K. J. and Capelson, M. (2017). Metazoan Nuclear Pores Provide a Scaffold for Poised Genes and Mediate Induced Enhancer-Promoter Contacts. Mol Cell 66, 63–76 e6.

Perkins, N. D. (2006). Post-translational modifications regulating the activity and function of the nuclear factor kappa B pathway. Oncogene 25, 6717–30.

Radhakrishnan, S. K. and Kamalakaran, S. (2006). Pro-apoptotic role of NF-kappaB: implications for cancer therapy. Biochim Biophys Acta 1766, 53–62.

Rasala, B. A., Orjalo, A. V., Shen, Z., Briggs, S. and Forbes, D. J. (2006). ELYS is a dual nucleoporin/kinetochore protein required for nuclear pore assembly and proper cell division. Proc Natl Acaď Sci U S A 103, 17801–6.

Rasala, B. A., Ramos, C., Harel, A. and Forbes, D. J. (2008). Capture of AT-rich chromatin by ELYS recruits POM121 and NDC1 to initiate nuclear pore assembly. Mol Biol Cell 19, 3982–96.

Riemer, D., Stuurman, N., Berrios, M., Hunter, C., Fisher, P. A. and Weber, K.(1995). Expression of Drosophila lamin C is developmentally regulated: analogies with vertebrate A-type lamins. J Cell Sci 108 (Pt 10), 3189–98.

Rushlow, C. A., Han, K., Manley, J. L. and Levine, M. (1989). The graded distribution of the dorsal morphogen is initiated by selective nuclear transport in Drosophila. Cell 59, 1165–77.

Ryan, K. M., Ernst, M. K., Rice, N. R. and Vousden, K.H. (2000). Role of NF-kappaB in p53-mediated programmed cell death. Nature 404, 892–7.

Sarkissian, T., Timmons, A., Arya, R., Abdelwahid, E. and White, K. (2014). Detecting apoptosis in Drosophila tissues and cells. Methoďs 68, 89–96.

Schultz, J., Milpetz, F., Bork, P. and Ponting, C. P. (1998). SMART, a simple modular architecture research tool: identification of signaling domains. Proc Natl Acaď Sci US A 95, 5857–64.

Silverman, N. and Maniatis, T. (2001). NF-kappaB signaling pathways in mammalian and insect innate immunity. Genes Dev 15, 2321–42.

Song, Z., McCall, K. and Steller, H. (1997). DCP-1, a Drosophila cell death protease essential for development. Science 275, 536–40.

Uv, A. E., Roth, P., Xylourgidis, N., Wickberg, A., Cantera, R. and Samakovlis, C. (2000). members only encodes a Drosophila nucleoporin required for rel protein import and immune response activation. Genes Dev 14, 1945–57.

Velentzas, P. D., Velentzas, A. D., Pantazi, A. D., Mpakou, V. E., Zervas, C. G., Papassideri, I. S. and Stravopodis, D. J. (2013). Proteasome, but not autophagy, disruption results in severe eye and wing dysmorphia: a subunit-and regulator-dependent process in Drosophila. PLoS One 8, e80530.

Vermaak, D., Hayden, H. S. and Henikoff, S. (2002). Centromere targeting element within the histone fold domain of Cid. Mol Cell Biol 22, 7553–61.

Vollmer, B., Lorenz, M., Moreno-Andres, D., Bodenhofer, M., De Magistris, P., Astrinidis, S. A., Schooley, A., Flotenmeyer, M., Leptihn, S. and Antonin, W. (2015). Nup153 Recruits the Nup107-160 Complex to the Inner Nuclear Membrane for Interphasic Nuclear Pore Complex Assembly. Dev Cell 33, 717–28.

Wagner, N., Weber, D., Seitz, S. and Krohne, G. (2004). The lamin B receptor of Drosophila melanogaster. J Cell Sci 117, 2015–28.

Waterhouse, A. M., Procter, J. B., Martin, D. M., Clamp, M. and Barton, G. J. (2009). Jalview Version 2--a multiple sequence alignment editor and analysis workbench. Bioinformatics 25, 1189–91.

Whalen, A. M. and Steward, R. (1993). Dissociation of the dorsal-cactus complex and phosphorylation of the dorsal protein correlate with the nuclear localization of dorsal. J Cell Biol 123, 523–34.

Wolff, T. (2011). Preparation of Drosophila eye specimens for scanning electron microscopy. Cold Spring Harb Protoc 2011, 1383–5.

Xu, L., Kang, Y., Col, S. and Massague, J. (2002). Smad2 nucleocytoplasmic shuttling by nucleoporins CAN/Nup214 and Nup153 feeds TGFbeta signaling complexes in the cytoplasm and nucleus. Mol Cell 10, 271–82.

Yang, X., Gu, Q., Lin, L., Li, S., Zhong, S., Li, Q. and Cui, Z. (2015). Nucleoporin 62-like protein activates canonical Wnt signaling through facilitating the nuclear import of beta-catenin in zebrafish. Mol Cell Biol 35, 1110–24.

Yokoyama, H., Koch, B., Walczak, R., Ciray-Duygu, F., Gonzalez-Sanchez, J. C., Devos, D. P., Mattaj, I. W. and Gruss, O. J. (2014). The nucleoporin MEL-28 promotes RanGTP-dependent gamma-tubulin recruitment and microtubule nucleation in mitotic spindle formation. Nat Commun 5, 3270.

Zuccolo, M., Alves, A., Galy, V., Bolhy, S., Formstecher, E., Racine, V., Sibarita, J. B., Fukagawa, T., Shiekhattar, R., Yen, T. et al. (2007). The human Nup107–160 nuclear pore subcomplex contributes to proper kinetochore functions. EMBO J 26, 1853–64.

